# Genetic compensation in β-actin mutants occurs independently of mutations that destabilize β-actin mRNA

**DOI:** 10.64898/2026.04.21.719943

**Authors:** Harleen Saini, Jiuchun Zhang, Hiba Dardari, Danesh Moazed

## Abstract

Proper maintenance of gene expression in response to mutations or environmental fluctuations is critical for cell development and survival. Recently, a novel genetic compensation mechanism was described wherein mutant mRNA decay triggers increased transcription of paralogous genes. This effect was reported for several genes, including β-actin (*Actb*) in mouse embryonic stem cells, where *Actb* mRNA with a premature termination codon enhances transcription of its paralog, γ-actin (*Actg1*), and partially rescues cytoskeletal defects. Here we show that, in both mouse and human embryonic stem cells, mutations in the *ACTB* gene, regardless of mutant mRNA expression, trigger genetic compensation. Furthermore, transgenic expression of mutant *ACTB* mRNA with a premature stop codon fails to induce genetic compensation. Depletion of the SRF or MRTF-A transcription factors, which are known to increase *ACTB* transcription in response to low ACTB protein levels, diminishes the genetic compensation response in *ACTB* mutants. These results suggest that genetic compensation in *ACTB* mutants is primarily mediated by a transcriptional feedback loop via SRF/MRTF-A, independent of the expression or degradation of mutant *ACTB* mRNA.

## Introduction

Genetic compensation refers to the ability of functionally redundant or paralogous genes to fully or partially compensate for each other’s loss (*1, 2*). When a gene is disrupted, compensation can be achieved by increasing the effective output of a paralogous gene through a variety of mechanisms – by increasing its transcription to make more mRNA, stabilizing its mRNA to make more protein, or simply stabilizing paralogous proteins to reach an appropriate steady-state expression level. In some cases, these changes are triggered by a loss or reduction of the mutant protein, which can activate transcription factor–based feedback loops (*3*), relieve transcriptional/translational repression (*3*), or reshape protein complex stoichiometry so that the paralogous protein is preferentially stabilized (*4*). Recent studies suggest a novel mechanism of genetic compensation triggered by mutant mRNA degradation and independent of protein loss (*5–10*).

mRNA degradation-dependent genetic compensation, also called transcriptional adaptation (*2*) or nonsense-induced transcriptional compensation (*11*), was first described for *egfl7* and *vegfaa* mutations in zebrafish (*5*). For *egfl7*, a vascular development gene, translation-blocking morpholinos targeting the wild-type mRNA caused clear vascular defects, but TALEN-generated loss-of-function alleles that produced a mutant *egfl7* mRNA (hereafter referred to as “gene-mutants”) showed little or no phenotype. The lack of mutant phenotype was attributed to rescue by increased expression of the related emilin family genes specifically in gene-mutant zebrafish, but not after knockdown by morpholinos or transcriptional inhibition by CRISPR interference (*5*). This compensatory response required components of the nonsense-mediated decay (NMD) pathway (*6*). Thus, genetic compensation appeared to require that mutant mRNA was produced and degraded. A similar phenomenon was reported for the calpain gene family, where morpholinos against *capn3a* caused liver defects but gene mutations that produced mutant mRNA did not, owing to increased expression of *capn3a* paralogs only in the gene-mutant zebrafish (*7*). Moreover, a premature termination codon (PTC)-containing *capn3a* transgene was sufficient to induce increased expression of its paralogs (*7*). In this case, genetic compensation specifically required the NMD factor Upf3a, but not the related NMD components Upf1, Upf2, or Upf3b (*7, 12*). Together, these observations led to the proposal that mutant mRNA degradation by NMD can actively increase transcription of related genes, providing a compensatory response that rescues protein loss. Similar genetic compensation responses have since been reported in nematodes (*8, 9, 13*), mouse (*6, 14*), and human (*10*) cell lines. It has been suggested that mutant mRNA or its fragments could directly influence transcription at paralogous genes in a sequence-specific manner, thereby maintaining both protein and mRNA homeostasis across gene families (*6, 7, 10, 13, 15*).

In mammalian cells, an mRNA degradation-dependent genetic compensation response has been described for the actin, kindlin (fermt), and rel gene families in mouse cell lines, and more recently for dystrophin in human cells (*6, 10, 14*). In the case of β-actin (*Actb*), a 4-nucleotide deletion (Δ4nt) in the coding sequence (CDS), which causes a frameshift and leads to a premature translation termination codon (PTC), was observed to result in enhanced transcription of its paralog, γ-actin (*Actg1*), whereas deleting the entire coding sequence of *Actb* (ΔCDS) failed to enhance *Actg1* transcription (*6*). Similarly, a deletion in the last coding exon of *Actg1*, which includes the canonical translation stop codon and is thought to undergo nonstop mRNA decay, caused increased transcription of its paralog *Actg2*, but a larger gene deletion failed to elicit this response (*6, 14*). Knocking down NMD factors, specifically eRF1 (ETF1), SMG6 or XRN1, diminished the genetic compensation response in the *Actb-Δ4nt* mutant (*6*). Thus, genetic compensation in actin mutants was concluded to require mutant mRNA degradation that directly feeds back to boost transcription of the paralogous gene(s), independently of protein loss (*6, 14, 15*). Additional experiments using heterozygous mutants and transfection of inherently unstable, uncapped mRNA provided evidence that the mutant mRNA can exert a dominant effect in genetic compensation (*6, 14*).

Actins are among the most highly conserved proteins, and actin-like proteins are found in all domains of life (*16*). In eukaryotes, filamentous actin plays central roles in cell motility, polarity, cytoskeletal organization, intracellular trafficking, muscle contraction, endocytosis, and signal transduction, while monomeric actin is a subunit of complexes that regulate actin filament assembly and chromatin remodeling, and other nuclear complexes (*17, 18*). Loss of actin expression or changes in actin protein levels are therefore likely to affect numerous essential cellular pathways. In humans and mice, the actin gene family comprises at least six paralogs with highly similar protein sequences (e.g. Actb and Actg1 differ by only four amino acids in their N-termini) but show functional divergence and tissue-specific expression (*19–21*). Whereas Actb and Actg1 are almost ubiquitously expressed, Acta1 is predominantly expressed in skeletal muscle, Actc1 in cardiac tissue, and Acta2 and Actg2 in smooth muscle. Several studies have noted that mutations in the *Actb* gene result in increased expression (RNA or protein) of actin paralogs (*Actg1*, *Acta1*, *Acta2*) (*22–27*).

Actin homeostasis in wild-type cells is exquisitely maintained by a protein-based feedback loop driven by the transcription cofactor MRTF-A, which binds monomeric globular actin (G-actin) (*3, 28–32*). When G-actin is depleted, either through enhanced polymerization into filamentous actin (F-actin) or, as in the case of *ACTB* mutations, an overall drop in actin protein, MRTF-A is released and associates with the transcription factor SRF to enhance the transcription of *Actb* and other cytoskeletal genes (*33*). However, the extent to which this protein-sensing mechanism contributes to the genetic compensation response at actin paralogs (*26, 30–33*) and how it relates to the recently described mRNA-dependent transcriptional adaptation mechanism (*6, 14*) remains unclear.

To investigate the mechanism of mutant mRNA-dependent genetic compensation and its relationship to actin protein feedback mechanism, we used previously described *Actb* mutants in mouse embryonic stem cells (mESCs) (*6*) and new *ACTB* mutants in human embryonic stem cells (hESCs) to specifically assess the role of mutant mRNA in the compensation response. We further tested whether mutant *ACTB* mRNA is sufficient to induce compensation using inducible transgenes and assessed the requirement for SRF/MRTF-A. Additionally, we examined previously described *Kindlin2* (*Fermt2*) mutants (*6*) as an alternative locus to assess the generality of mutant-mRNA–triggered compensation. Our results show that, in *ACTB* mutant hESCs, genetic compensation is mediated by an SRF/MRTF-A–dependent protein feedback response and does not require mutant mRNA decay. Moreover, in hESCs and mouse kidney fibroblasts, heterozygous *Kindlin2* mutations fail to trigger genetic compensation at the paralog *Kindlin1*.

## Results

### Genetic compensation in β-actin mutants occurs independently of mutant mRNA expression in human and mouse embryonic stem cells

To dissect the mechanism of genetic compensation, we generated two types of heterozygous *ACTB* mutants in human embryonic stem cells (hESCs). The first is a 4-nucleotide deletion (*ACTB-Δ4nt^+/-^*) that causes a frameshift resulting in a premature termination codon (PTC), mimicking the previously described mouse *Actb* mutation that triggers mutant mRNA-dependent genetic compensation (*6*). The second is a 3.5-kb genomic deletion that removes the entire *ACTB* gene (*ACTB-Δgene^+/-^*) (**Fig. 1A and Fig. S1A**). Both mutants eliminate functional β-actin protein from one allele, but only *ACTB-Δ4nt^+/-^* is expected to express a mutant mRNA containing a PTC. Surprisingly, RT-qPCR showed increased mRNA expression of *ACTB* paralogs (*ACTA1* and *ACTG1*) in both mutant lines, indicating that genetic compensation can occur even in the absence of a mutant *ACTB* mRNA from the edited allele (**Fig. 1B**). Notably, total *ACTB* mRNA levels were not reduced in the *ACTB-Δgene^+/-^* heterozygous mutant, likely due to increased expression from the remaining wild-type allele. Thus, in hESCs, loss of one functional *ACTB* allele is sufficient to trigger genetic compensation independently of mutant mRNA expression.

**Fig. 1.**
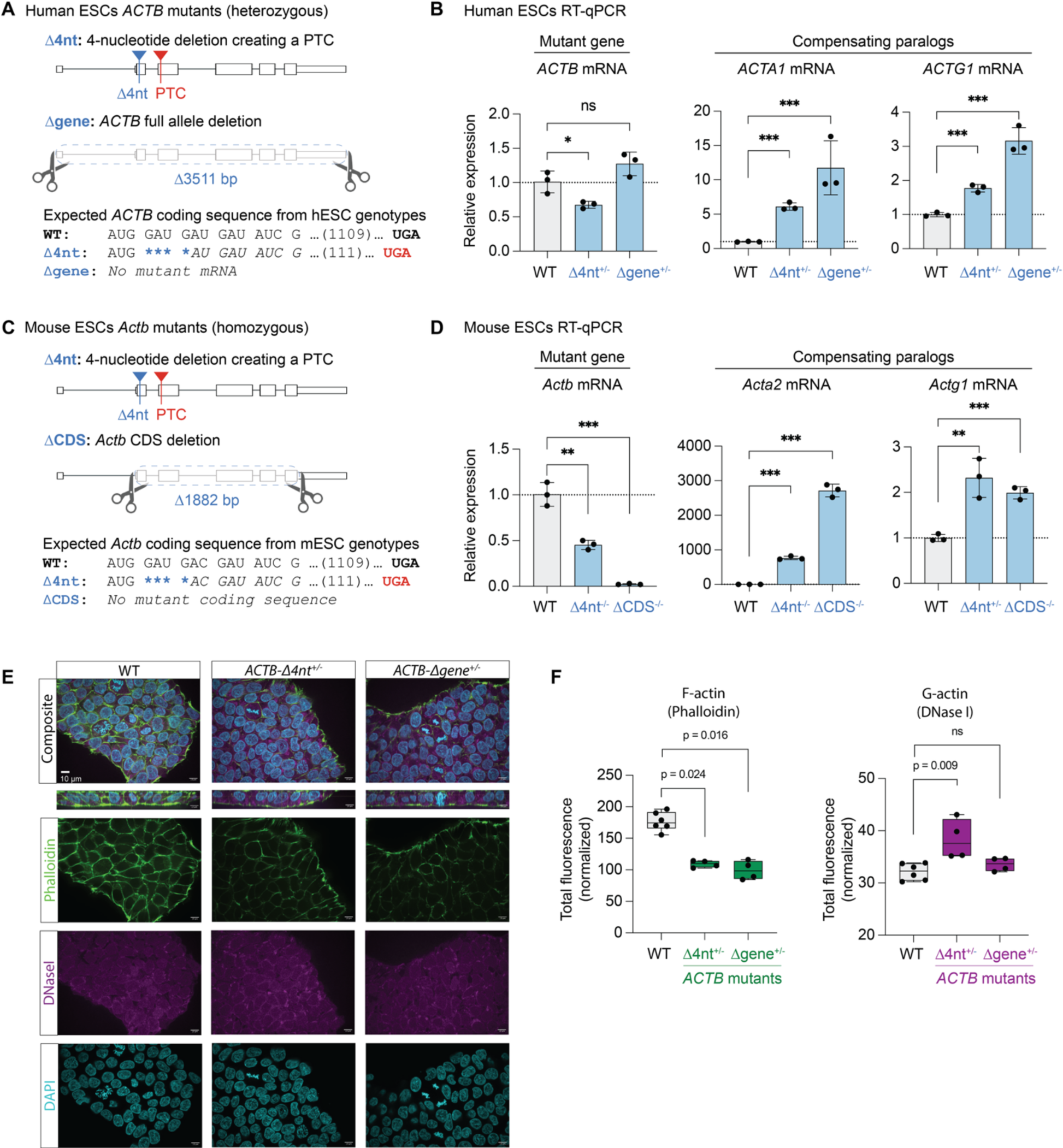
Genetic compensation in β-actin mutants occurs independently of mutant mRNA expression in human and mouse embryonic stem cells. **(A)** Schematic of heterozygous *ACTB* mutant alleles in hESCs, with corresponding expected *ACTB* mRNA coding sequence shown below the schematic. The PTC (premature termination codon) in Δ4nt is labeled in red; the Δgene allele deletes the *ACTB* transcribed region and is not expected to produce an *ACTB* transcript from the deleted allele. **(B)** RT-qPCR analysis of mRNA expression of *ACTB* and its paralogs (*ACTA1* and *ACTG1*) in hESCs. Relative expression was calculated using the 2-ΔΔCt method, normalized to *PGK1* mRNA and wild-type (WT) cells. Error bars denote standard deviation; p-values were calculated with a two-tailed t-test on ΔCt values (gene-of-interest – PGK1). p-value < 0.05 (*), < 0.01 (**), <0.001 (***). **(C)** Schematic of homozygous *Actb* mutations in mESCs with corresponding expected *Actb* mRNA coding sequence (CDS, coding sequence). The ΔCDS allele removes the *Actb* CDS (but not flanking transcribed regions). **(D)** RT-qPCR analysis of *Actb* and its paralogs (*Acta2* and *Actg1*) mRNA expression in mESCs. Relative expression was calculated using the 2-ΔΔCt method, normalized to *Gapdh* mRNA and WT cells. Error bars and p-values were calculated as in (B). **(E)** Representative immunofluorescence images of F-actin and G-actin in hESCs. From top to bottom: Composite image showing F-actin (phalloidin, green), G-actin (DNaseI, purple), and DNA (DAPI, cyan); orthogonal section through the z-stack, to show cell depth (thin plane); individual channels. Each column (left to right) corresponds to the labeled genotype. Scale bar = 10 µm. **(F)** Quantification of F-actin (phalloidin, left) and G-actin (DNase I, right) fluorescence, summed across the z-stack and normalized to the total area covered by cells in each image. Each dot in the box plot represents a distinct image containing multiple cells.

Because the hESC *ACTB* mutants showed compensation even without mutant mRNA, we next examined previously described mouse embryonic stem cell (mESC) lines in which *Actb* mutations were reported to trigger genetic compensation only when a mutant mRNA was present, suggesting that the mutant mRNA was necessary for the response (*6*). We used *Actb* mutant mESC lines with either a 4-nucleotide deletion in the *Actb* coding sequence that produces a PTC-containing mutant transcript (*Actb-Δ4nt^-/-^*), or a deletion of the entire coding sequence (*Actb-ΔCDS^-/-^*), such that no coding sequence-containing mutant mRNA is produced (**Fig. 1C and Fig. S1B**), to test effects on expression of paralogs. Contrary to previous findings, we observed increased expression of *Actg1* in both mESC mutants (**Fig. 1D**). Similar to *Actg1*, *Acta2* mRNA was also significantly increased in both mESC mutants. These results indicate that genetic compensation in both mouse and human ESCs can occur through an mRNA-independent mechanism. The reason for the discrepancy between our results and previous findings (*6*) is unclear but may stem from differences in clonal state, culture conditions, and/or regulatory adaptation to reduced expression or loss of a critical protein.

We then tested whether genetic compensation rescues cytoskeletal defects by assessing filamentous (F-actin) and the DNase I-accessible monomeric actin pool (G-actin). Both hESC mutant lines exhibited a similar and significant reduction in F-actin (phalloidin staining; **Fig. 1E, F**), indicating that the compensatory transcriptional response is not sufficient to fully restore actin filament organization, and that expression of mutant *ACTB* mRNA in *ACTB-Δ4nt^+/-^*conferred no detectable improvement. DNase I staining was increased only in *ACTB-Δ4nt^+/-^*, suggesting an allele-specific effect on the DNase I-accessible monomer pool (**Fig. 1E, F**). The mechanistic basis of reduced F-actin in *ACTB* mutants despite compensatory changes in actin transcripts and elevated or unchanged G-actin remains unclear, but may result from altered actin isoform composition (*25, 34*).

Together, these results demonstrate that genetic compensation in both human and mouse ESCs occurs independently of mutant mRNA expression and does not restore normal actin filament architecture, even in heterozygous mutants.

### Transgenic expression of mutant *ACTB* mRNA is insufficient to trigger genetic compensation

To directly test whether mutant mRNA is sufficient to trigger a genetic compensation response, we engineered doxycycline-inducible transgenes encoding either wild-type or nonsense mutant *ACTB* (*ACTB-insPTC*) mRNA and integrated them into the genome of wild-type hESCs (**Fig. 2A**). Upon transcription induction by doxycycline (dox), wild-type *ACTB* expression led to a significant decrease in *ACTA1* and *ACTG1* mRNA, as measured by RT-qPCR (**Fig. 2B, C**). The decrease in *ACTA1* and *ACTG1* expression upon ACTB overexpression is consistent with a protein feedback mechanism in which G-actin sequesters the MRTF-A transcription cofactor in the cytoplasm, thereby reducing SRF-dependent transcription of actin family genes (*28, 30–33*). Consistent with this hypothesis, expression of a nonsense mutant *ACTB* mRNA from an *ACTB* transgene containing two adjacent PTCs in the CDS (*ACTB-insPTC* transgene), and thus expected to produce no full-length β-actin protein, did not alter paralog expression (*ACTA1* or *ACTG1*), indicating that mutant *ACTB* mRNA alone is insufficient under these conditions to engage either MRTF-A/SRF feedback or transcriptional adaptation (**Fig. 2D, E**). Similar results were obtained in HEK293FT cells (**Fig. S2**). These results further support the model whereby genetic compensation in *ACTB* mutants is primarily a response to protein-level changes, rather than to mutant mRNA expression.

**Fig. 2.**
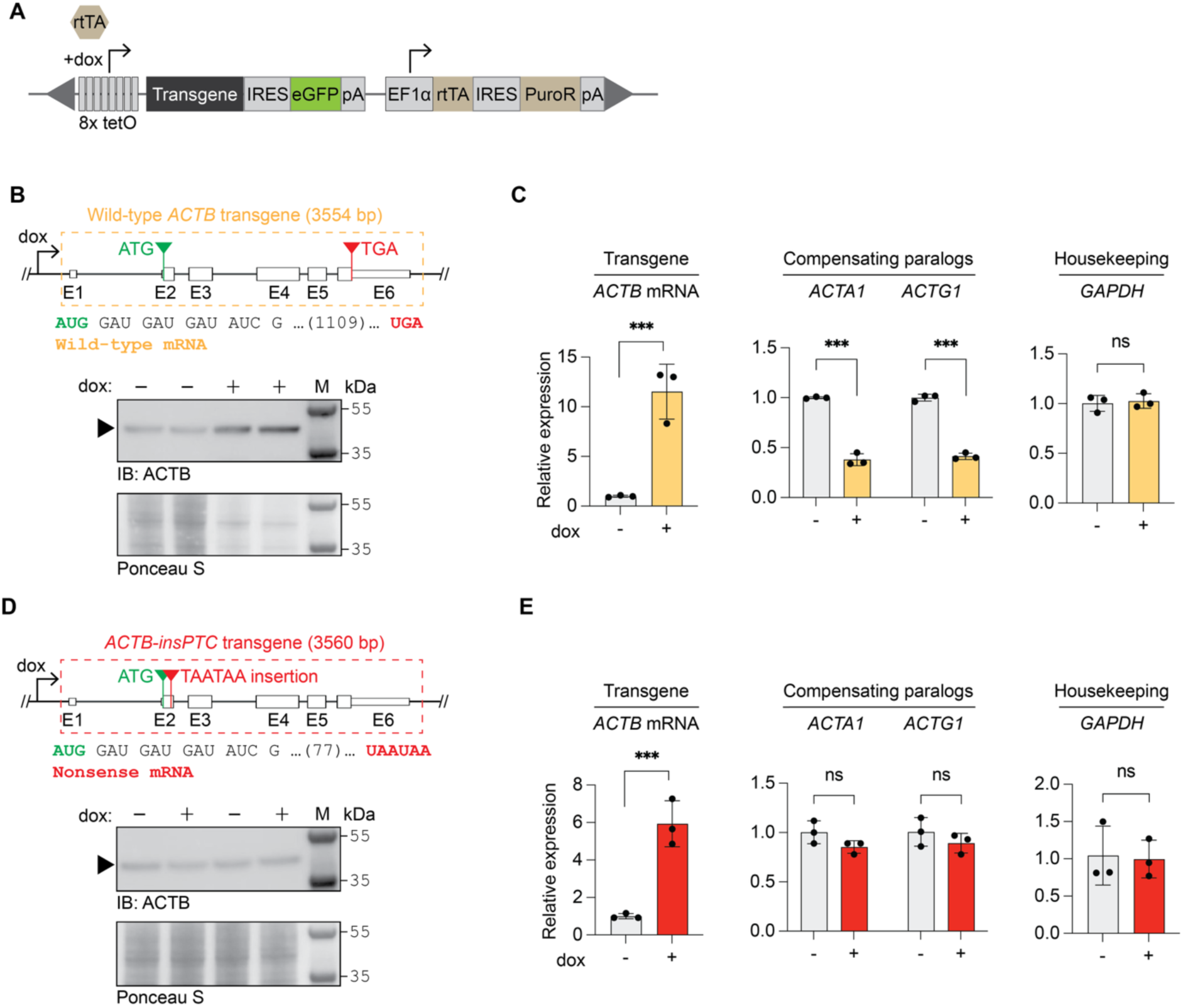
Transgenic expression of mutant *ACTB* mRNA does not induce genetic compensation. **(A)** Schematic of the doxycycline-inducible construct used for transgene integration and expression. An 8x tetO sequence is positioned upstream of the transgene, IRES, and eGFP, followed by a polyA signal. An EF1a promoter drives rtTA and puromycin resistance expression. The entire cassette is flanked by inverted terminal repeats (grey triangles) for piggyBac transposase-mediated integration. Arrows mark transcription start sites for dox-inducible (8x tetO) and constitutive (EF1a) promoters. Abbreviations: rtTA, reverse tetracycline transactivator; 8x tetO, eight tandem tet operator repeats; IRES, internal ribosome entry site; eGFP, enhanced green fluorescent protein; EF1a, rat elongation factor 1α promoter; pA, polyadenylation sequence; PuroR, puromycin resistance gene; dox, doxycycline. Schematic adapted from Kim et al., 2011. **(B)** (Top) Schematic of the wild-type *ACTB* transgene. The sequence corresponds to chr7:5527048–5530601 (hg38). Exons (E1–E6) are represented as rectangles (coding exons are shown as taller boxes than untranslated regions [UTRs]), with introns indicated by connecting lines. The start codon (ATG) is highlighted in green, and the stop codon (TGA) in red. The start and end of the wild-type *ACTB* mRNA sequence is displayed beneath the gene schematic. (Bottom) Immunoblot for ACTB protein expression in cells before and after 24 hours of doxycycline (dox) treatment. Black arrowhead points to band corresponding to ACTB. ‘M’ indicates the protein molecular weight marker. Ponceau S staining of the membrane is shown as a loading control. **(C)** RT-qPCR analysis of *ACTB*, its paralogs (*ACTA1* and *ACTG1*), and a housekeeping gene (*GAPDH*) mRNA expression before (–) or after (+) 24 hours of wild-type *ACTB* transgene induction by doxycycline. Relative expression was calculated using the 2-ΔΔCt method, normalized to *PGK1* mRNA. Error bars denote standard deviation. p-values were calculated with a two-tailed t-test on ΔCt values (gene-of-interest-PGK1). p-value < 0.05 (*), < 0.01 (**), <0.001 (***). **(D, E)** Same as in (B, C) but for the nonsense mutant *ACTB-insPTC* transgene, which contains two adjacent PTCs highlighted in red.

### *ACTB* mutations increase SRF and SRF target gene expression in hESCs

To examine the global effects of *ACTB* mutations, we sequenced polyadenylated mRNA from wild-type hESCs and two *ACTB* mutant lines: the *ACTB-Δ4nt^+/-^* mutant and a newly designed *ACTB-insPTC^+/-^* allele, in which two adjacent stop codons (TAA TAA) were inserted in exon 2 of *ACTB* (**Fig. S1A, S3A**) and are expected to induce nonsense-mediated decay of the mRNA.

An allele-specific single nucleotide polymorphism (SNP) in the *ACTB* 3′ UTR allowed us to directly assess whether the mutant allele is selectively degraded (**Fig. 3C**). Whereas wild-type cells showed balanced biallelic *ACTB* mRNA expression (45%:55%), the mutants had skewed allelic expression (95%:5% in *ACTB-Δ4nt^+/-^* and 64%:36% in *ACTB-insPTC^+/-^*), consistent with preferential loss of mutant mRNA, which was more pronounced in the *ACTB-Δ4nt^+/-^* mutant cells (**Fig. 3D**).

**Fig. 3.**
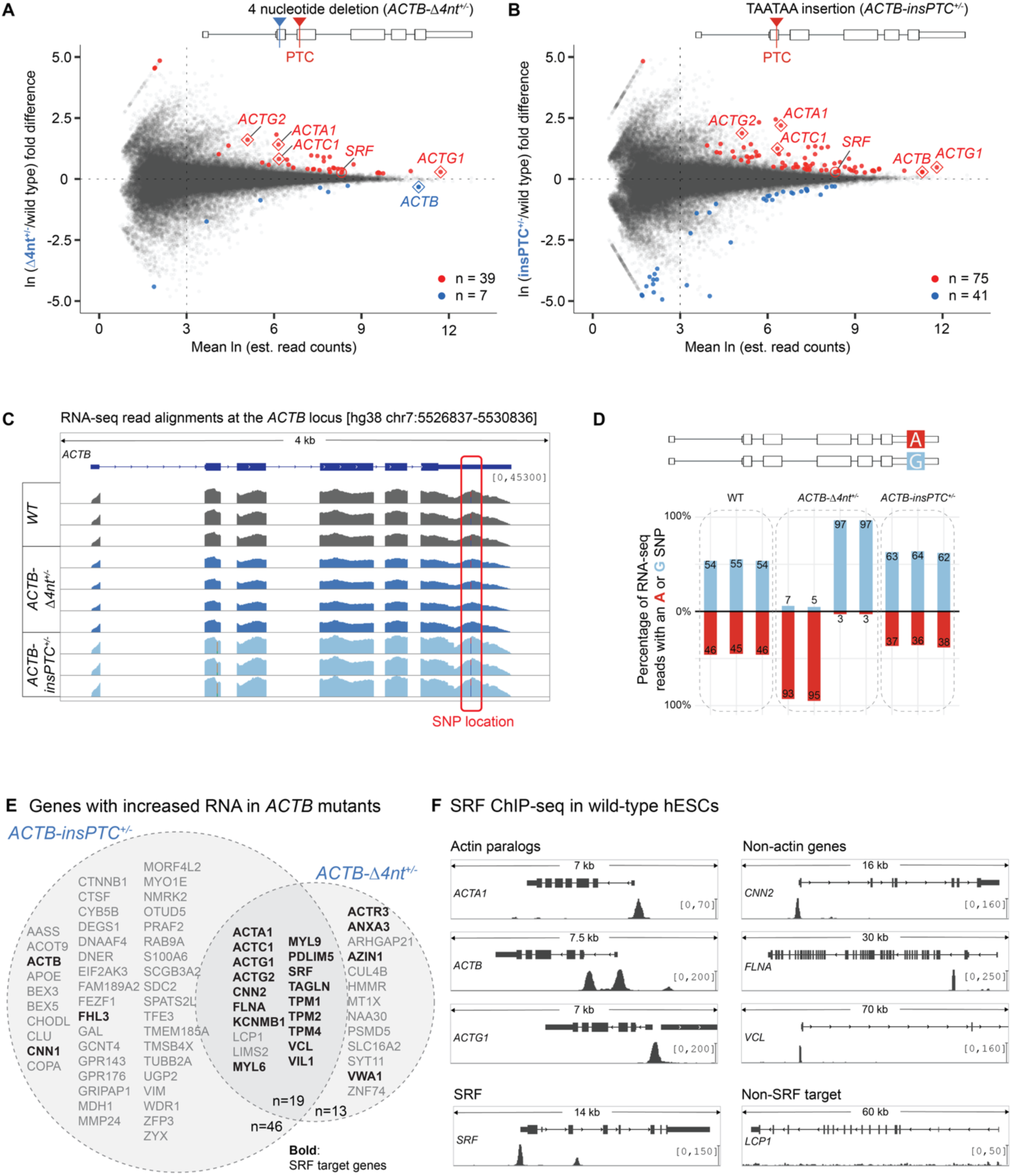
Increased expression of *SRF* and SRF-target genes in *ACTB* mutants. **(A, B)** Scatter plot showing RNA-seq quantification, comparing log_e_ mean read counts and log_e_ fold difference in expression in *ACTB-Δ4nt^+/-^* (A), or *ACTB-insPTC^+/-^* (B) mutant versus wild-type hESCs per annotated mRNA isoform. Transcripts with significant changes in abundance (q-value < 0.01) are labeled in red (increased expression) or blue (decreased expression). Actin paralogs and *SRF* transcription factor are annotated. Schematic of the corresponding *ACTB* mutant allele is shown above the scatter plot. **(C)** Integrative Genomics Viewer (IGV) screenshot showing read coverage across a 4-kb region of the *ACTB* locus for wild type (WT, gray), *ACTB-Δ4nt^+/−^* (dark blue), and *ACTB-insPTC^+/−^* (light blue) hESCs. Each track represents an RNA-seq biological replicate (independent clones). Coverage tracks were generated from non-normalized BAM files and group-scaled across all samples to a common y-axis range of 0–45,300, corresponding to the absolute read counts at each position. The position of the heterozygous SNP in the *ACTB* 3′ UTR is indicated; individual reads overlapping the SNP are colored red or blue according to the nucleotide present at that position. **(D)** Allele-specific *ACTB* expression in mutant hESCs. Bar plots showing the percentage of RNA-seq reads carrying either the G (blue) or A (red) nucleotide at the heterozygous SNP in the *ACTB* 3′ UTR for each indicated clone. A schematic of the SNP position and allele identities is shown above the plot. All samples have >16,000 reads overlapping the SNP, enabling robust estimation of allelic bias in *ACTB* mRNA expression. **(E)** Venn diagram showing genes with increased RNA abundance in *ACTB* mutants. SRF target genes are highlighted in bold black font. **(F)** Genome browser views showing SRF occupancy by ChIP-seq in wild-type hESCs on a subset of genes with increased RNA expression in *ACTB* mutants. For each gene, the top track shows the gene annotation from GRCh38, the bottom track shows SRF ChIP-seq read coverage as fold change over control. The genome browser track was downloaded from ENCODE (experiment ENCSR000BIV, file ENCFF23158KUZ) and is displayed on IGV.

Extending earlier observations in mESCs that *Actg1* expression is increased in *Actb* mutant cells (*6*), and consistent with our RT-qPCR data in Fig. 1A–D, RNA-seq analysis showed that all *ACTB* paralogs (*ACTA1*, *ACTC1*, *ACTG1*, and *ACTG2*) had significantly increased steady-state RNA abundance in *ACTB-Δ4nt^+/-^*and *ACTB-insPTC^+/-^* mutant hESCs (**Fig. 3A, B**). *ACTB-insPTC^+/-^*mutant ESCs also showed increased *ACTB* mRNA, consistent with feedback regulation at the *ACTB* locus (**Fig. 3B, C**). Preferential decay of the Δ4nt or insPTC transcript is expected to bias steady-state RNA toward the wild-type allele (**Fig. 3D**). Of note, among the genes with significantly increased RNA expression was *SRF*, a transcription factor important for maintaining ACTB homeostasis, along with several known targets of SRF (**Fig. 3C**). Of the 19 genes upregulated in both mutants, 17 genes, including *SRF* itself, had SRF ChIP-seq occupancy at their promoters in published hESC datasets (ENCODE (*35*)), suggesting that they may be directly activated by SRF in the mutant hESCs (**Fig. 3E, F**). Consistent with the hESC results, both mESC *Actb* mutant lines also showed elevated *Srf* mRNA, and the Srf target gene *Vcl* was also increased, as assessed by RT-qPCR (**Fig. S3B, S3C**). These results suggest that SRF may serve as a key effector in regulating the transcription of the actin gene family and other cytoskeletal genes, leading to the genetic compensation response in *ACTB* mutants in both human and mouse ESCs.

### SRF is required for compensatory actin paralog induction in *ACTB* and *ACTG1* mutants

Compensation for actin loss may occur through multiple pathways, including SRF/MRTF-A–dependent feedback that maintains actin/cytoskeletal gene expression (*3*), and mutant mRNA–dependent transcriptional adaptation (*6*), which was recently reported to require ILF3 (*36*). To directly test the role of SRF/MRTF-A in *ACTB* mutants and determine whether mutants with or without mutant mRNA expression use different transcriptional activation mechanisms, we knocked down SRF or MRTF-A using siRNAs. Genetic compensation was abolished upon depletion of SRF/MRTF-A regardless of mutant mRNA expression, suggesting a key role for MRTF-A/SRF in genetic compensation that is independent of mutant mRNA expression (**Fig. 4A, B; Fig. S3E–G** for knockdown efficiencies).

**Fig. 4.**
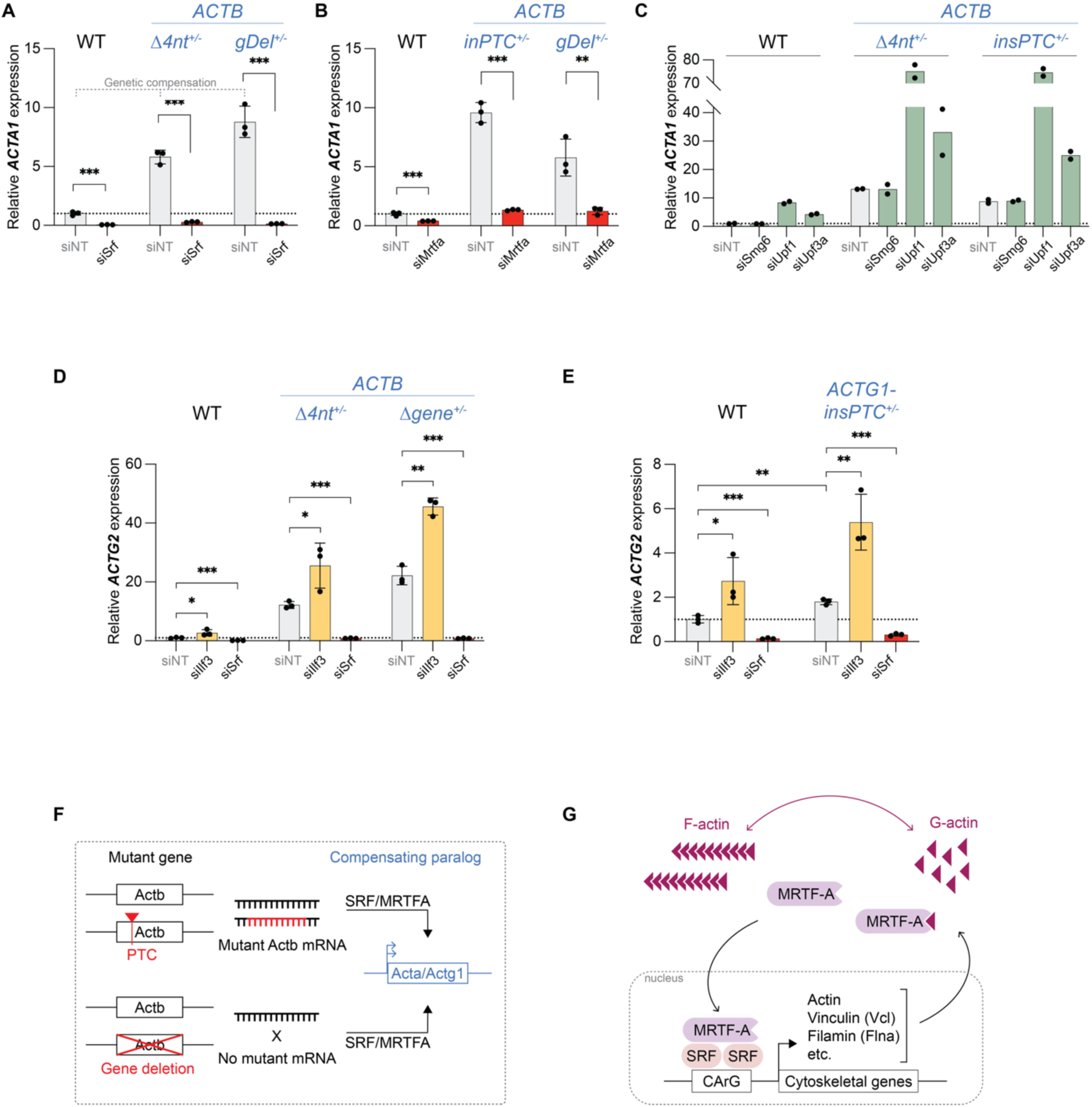
Genetic compensation in *ACTB* and *ACTG1* mutants requires SRF/MRTF-A and is independent of NMD factors and ILF3. **(A, B)** RT-qPCR analysis of *ACTA1* mRNA expression in wild type or *ACTB* mutants after siRNA-mediated knockdown of SRF (A) or MRTF-A (B). The genetic compensation response can be seen by comparing the siNT condition across samples. Relative expression was calculated using the 2-ΔΔCt method, normalized to *PGK1* mRNA and siNT control in wild type cells. Error bars denote standard deviation. p-values were calculated with a two-tailed t-test on ΔCt values (gene-of-interest – PGK1). p-value < 0.05 (*), < 0.01 (**), <0.001 (***). NT, Non-targeting control siRNA. **(C)** RT-qPCR analysis of *ACTA1* mRNA expression after knockdown of NMD factors, SMG6, UPF1, or UPF3A, by siRNA. Relative expression, error bars, and p-values were calculated the same way as in (A, B) above. **(D, E)** RT-qPCR analysis of *ACTG2* mRNA expression after knockdown of SRF or ILF3 in *ACTB* (D) or *ACTG1* (E) mutants. **(F)** Model for SRF/MRTF-A–dependent genetic compensation in *ACTB* mutants. Schematic illustrating heterozygous *ACTB* alleles carrying either a premature termination codon (PTC; top) or a gene deletion (bottom). In the PTC allele, mutant *ACTB* mRNA is transcribed and degraded, whereas in the gene-deletion allele no transcript is produced. In both cases, reduced ACTB protein leads to MRTF-A/SRF-dependent increased transcription of actin paralogs (e.g., *ACTA1*, *ACTG1*, *ACTG2*), indicating that genetic compensation depends on SRF/MRTF-A–mediated protein-sensing rather than the presence of mutant *ACTB* mRNA. **(G)** Model of MRTF-A/SRF-mediated feedback in ACTB overexpression versus mutant mRNA expression. MRTF-A partners with SRF transcription factor to drive the expression of cytoskeletal genes, including actins. When wild-type ACTB is overexpressed, increased G-actin sequesters MRTF-A in the cytoplasm and limits its nuclear localization, leading to reduced MRTF-A/SRF activity and decreased transcription of actin paralogs. In contrast, ectopic expression of nonsense mutant *ACTB* mRNA does not change ACTB protein levels, leaving MRTF-A/SRF activity at baseline and paralog expression unchanged. Thus, mutant *ACTB* mRNA alone is ineffective in inducing genetic compensation. Figure adapted from Pipes et al. 2006 (*38*).

To test whether NMD factors were also required for genetic compensation in these *ACTB* mutants, we knocked down key NMD factors: SMG6, UPF1, or UPF3A (**Fig. S3H–J**). While loss of SMG6 had no effect on *ACTA1* mRNA expression, UPF1 and UPF3A depletion resulted in increased expression of *ACTA1* in both wild-type and *ACTB* mutant hESCs, indicating that the NMD pathway is not required for genetic compensation at actin paralogs (**Fig. 4C and Fig. S3D** for an alternative view of normalized expression change). This is consistent with prior RNA-seq analysis of UPF1-depleted hESCs, in which several actin isoform transcripts (including *ACTA1*, *ACTC1*, and *ACTG2*) increased upon UPF1 knockdown (Lou et al., 2016; Table S1) (*37*).

A recent study identified Ilf3 as an effector necessary for increased expression of the *Actg2* gene in mouse embryonic fibroblasts that express an unstable/mutant *Actg1* mRNA (*36*). In the same study, a CRISPR screen for *Actg2* induction also identified SRF (broadly) and MRTF-A (*Actg1*-mutant-specific) as top hits (**Fig. S4A, B**). Our results showed that the expression of *ACTG2* is also robustly increased in *ACTB* mutant hESCs (**Fig. 3A, B**). We therefore tested whether depletion of ILF3 impaired genetic compensation via *ACTG2* upregulation in the *ACTB-Δ4nt^+/-^* and *ACTB-Δgene^+/-^* hESCs by knocking down either ILF3 or SRF using siRNAs (**Fig. S3K, L**). Depletion of ILF3 resulted in increased, rather than decreased, *ACTG2* expression in both *ACTB* mutant hESCs (**Fig. 4D**), indicating that ILF3 was not required for the observed genetic compensation. ILF3 was also not required for increased expression of *ACTG2* in *ACTG1-insPTC^+/-^* mutant hESCs (**Fig. 4E, Fig. S1A, Fig. S4C-F**). As a control and consistent with the results in Fig. 3, depletion of SRF abolished *ACTG2* upregulation in both *ACTB* and *ACTG1* mutant hESCs (**Fig. 4D, E**). We conclude that ILF3-mediated transcriptional adaptation is not required for increased expression of *ACTG2* in *ACTB* or *ACTG1* mutant hESCs.

### Heterozygous *Kindlin2* mutations do not trigger genetic compensation in hESCs and mouse kidney fibroblasts

A previous study in mouse kidney fibroblasts (MKFs) reported that a ∼600 bp deletion in *Kindlin2* robustly increased transcription of the paralog *Kindlin1* (*Fermt1*) and interpreted this as mutant-mRNA–triggered compensation (*6*). We generated heterozygous *Kindlin2* mutations in hESCs and MKFs. Although *Kindlin2* mRNA level was reduced in both hESCs and MKF heterozygous mutants, *Kindlin1* expression was not significantly increased (**Fig. S5A, B**). In hESCs and MKFs, a nonsense or nonstop mutation in *Kindlin2* is therefore insufficient to induce genetic compensation by increasing the expression of its paralog (**Fig. 5C**).

## Discussion

Our findings demonstrate that increased expression of actin paralogs in human ESCs, resulting from mutations in the *ACTB* gene, is primarily driven by the loss of ACTB protein and occurs independently of mutations that produce unstable *ACTB* mRNA. We provide several lines of evidence that this genetic compensation is mediated by a protein sensing mechanism that requires the SRF transcription factor and its co-factor, MRTF-A, which have been shown to regulate the expression of actin and other cytoskeletal genes in response to changes in G-actin levels (*28–33, 38*). First, an *ACTB* full-locus gene deletion allele and *ACTB* mutant alleles with premature stop codons elicit a similar increase in actin paralog expression in both human and mouse ESCs. Second, this increased paralog expression is lost upon the depletion of either MRTF-A or SRF. Third, transgenic expression of nonsense mutant *ACTB* mRNA in cells carrying wild-type *ACTB* fails to increase the expression of *ACTB* paralogs, while the same nonsense mutation, when present at the endogenous locus, shows a robust compensatory response. Finally, consistent with negative feedback through an actin protein-sensitive pathway, overexpression of wild-type *ACTB* suppresses *ACTG1* expression. Below, we discuss the implication of these findings for mechanisms that maintain actin homeostasis via genetic compensation.

Actins are among the most highly expressed and conserved eukaryotic proteins and impact every aspect of cell biology (*16*). In addition to pathways that regulate actin polymerization dynamics, actin expression is regulated at the level of transcription (*3*). MRTF-A and SRF form a complex that activates the transcription of *ACTB* and its paralogs, and MRTF-A is sequestered in the cytoplasm by G-actin. Reduced G-actin levels are thought to increase free MRTF-A levels and allow its translocation into the nucleus leading to increased expression of actin paralogs and other cytoskeletal proteins. MRTF-A therefore serves as a sensor for actin protein levels. Our results suggest that this sensing mechanism is required for the upregulation of actin paralogs in cells that carry either a complete *ACTB* gene deletion or *ACTB* alleles that contain PTCs and produce unstable mRNA (**Fig. 4F**). In our experiments, ectopic expression of a PTC-bearing *ACTB* mRNA transgene in otherwise wild-type cells did not induce increased expression of actin paralogs. This, combined with the observation that PTC and full gene deletion alleles both increase paralog expression, suggests that the protein-sensing mechanism is the dominant mechanism in hESCs (**Fig. 4G**). More generally, these findings highlight that protein-sensing feedback can be sufficient to drive compensation within highly buffered gene networks such as the actin cytoskeleton (*3*).

Previous studies have shown that mRNA decay-dependent transcriptional adaptation contributes to genetic compensation in mRNA destabilizing mutations for multiple genes in zebrafish, mouse, and human cells (*5–10, 14*). The evidence for transcriptional adaptation in some of these cases is compelling, as, for example, multiple PTC-bearing mutations in the zebrafish *capn3a* (*7*) and the mouse *Actg1* (*6, 14*), but not their complete gene knockouts, lead to increased expression of their respective paralogs. From an evolutionary perspective, transcriptional adaptation provides a general and highly versatile mechanism that can increase the expression of paralogous genes, and even a remaining wild-type allele, in response to mutations that impair the function of any gene in the family and are linked to decay of mutated gene’s mRNA. The power of such RNA-based systems is exemplified by numerous examples of RNAi and CRISPR surveillance pathways. On the other hand, protein-sensing compensation mechanisms lack the generality of mRNA decay-dependent transcriptional adaptation and require the evolution of a specific sensor that mediates compensation for each gene family. Our findings on the mechanism of genetic compensation associated with actin and kindlin mutations in ESCs are therefore surprising and suggest that transcriptional adaptation may be gene- and/or cell type-sensitive.

## Acknowledgments

We thank Anna Kashina for advice on actin biology, and Sean Eddy, Fred Winston, Didier Stainier, Mohamed El-Brolosy, and Jonathan Weissman for feedback on the manuscript. We are grateful to Robi Mitra for providing the piggyBac hyPBase plasmid, and to Mohamed El-Brolosy, Zacharias Kontarakis, and Didier Stainier for sharing cell lines and protocols, and advice on culturing mESC and MKF lines. We acknowledge the Harvard Medical School Imaging Technology & Education (CITE) core for microscopy support and the Image Analysis Collaboratory (IAC) for guidance on image analysis. We thank the HMS Cell Biology Initiative for Genome Editing and Neurodegeneration for assistance with hESC culture and genome editing. We thank Yekaterina Shulgina and members of the Moazed laboratory, Roberto Sotomayor, Yizhe Zhang, Antonis Tatarakis, Juntao Yu, Ana Dorador, and Swapnil Parhad for helpful feedback.

## Funding

D.M. is an investigator of the Howard Hughes Medical Institute.

HHMI Helen Hay Whitney Fellowship (HS)

Cell Biology Education and Fellowship Fund (HS)

Harvard College Research Program Fellowship (HD)

Howard Hughes Medical Institute (DM)

## Author contributions

H.S. and D.M. conceived the study and designed the overall experimental strategy. hESC mutants were generated by J.Z. H.D. contributed to MKF *Kindlin2* mutant investigations. H.S. conducted all other experiments and analyzed the data with supervision from D.M. H.S. and D.M. wrote the manuscript with feedback from all authors.

## Competing interests

Authors declare that they have no competing interests.

## Data, code, and materials availability

All plasmids generated for this study are deposited to Addgene. RNA-seq data are available at NCBI GEO accession: GSE317480 (**reviewer token**: gpebeuywltwtvkn). Raw imaging data are available on Zenodo (https://doi.org/10.5281/zenodo.19556498). Image analysis code is available on GitHub (https://github.com/harleensaini/MoazedLab_hESC-actin-quantification). All other data, code, and materials used in the analysis are provided in the main text or supplementary materials.

## Materials and Methods

### Cell lines and culture conditions

All cell lines were maintained in humidified 37°C incubators with 5% CO2.

#### Human embryonic stem cells (hESCs)

H9 hESCs were obtained from WiCell. Cells were cultured on Matrigel-coated plates (Corning® Matrigel® GFR Basement Membrane Matrix, LDEV-free) and maintained in feeder-free E8 medium. Medium was changed daily, and cells were passaged every 2-3 days using 0.5 mM EDTA in 1x DPBS for dissociation. E8 medium contained DMEM/F-12 (Thermo Fisher Scientific: 11330032), sodium bicarbonate (Sigma: S3817), sodium selenite (Sigma: S5261), L-ascorbic acid 2-phosphate (Sigma: A8960), human insulin (Sigma: I9278), human holo transferrin (Sigma: T0665), recombinant human TGF-β1 (PeproTech: 100-21), and human FGF basic (PeproTech: 100-18B).

#### Mouse embryonic stem cells (mESCs)

Wild-type and *Actb* mutant mESCs were provided by Didier Stainier’s laboratory (Max Planck Institute for Heart and Lung Research in Bad Nauheim, Germany) (*6*). Cells were cultured following the protocol published in El-Brolosy et al. (*6*). Briefly, mESCs were cultured on 0.1% gelatin coated plates and maintained in DMEM (Invitrogen: 10313-039) supplemented with 15% embryonic stem cell FBS (Invitrogen: 10439001), 2 mM glutamine (Invitrogen: 25030081), 1% non-essential amino acids (Invitrogen: 11140050), 100 U/ml Penicillin-Streptomycin (Invitrogen: 15140-122), 0.1 mM β-mercaptoethanol, 1000 U/ml mLeukemia Inhibitory Factor (STEMCELL™ Technologies: 78056), 3 µM CHIR99021 (Sigma: SML1046), and 1 µM PD0325901 (Sigma: PZ0162). Cells were passaged every 2-3 days using Accutase® Cell Dissociation Reagent (Thermo Fisher Scientific: A1110501). mESC mutant genotypes were verified by PCR and sequencing the PCR product (see Supplementary Data S1 for genotyping primers).

#### Mouse kidney fibroblasts (MKFs) and HEK293FT cells

Parental and homozygous mutant MKFs were provided by Mohamed El-Brolosy and Didier Stainier (*6*). Cells were cultured following the protocol published in El-Brolosy et al. (*6*). MKFs were maintained in DMEM (Invitrogen: 11995073) with 10% fetal bovine serum (SeraPrime®: F31016-500) and passaged every 4-5 days using 0.05% Trypsin-EDTA (Thermo Fisher Scientific: 25300120) for dissociation. Genotypes were verified by PCR product size (See Supplementary Data S1 for genotyping primers).

HEK293FT cells (Invitrogen: R70007) were cultured using the same conditions as described for MKFs.

### Generation of mutant and transgenic cell lines

#### Endogenous gene editing in hESCs

HiFi Cas9 (R691A) expression plasmid was generated by site-directed mutagenesis of pET-Cas9-NLS-6xHis (Addgene: 62933 (*39*)). HiFi Cas9 protein was expressed in Rosetta™(DE3)pLysS Competent Cells (Novagen) and purified following established protocols (*40*). The AsCas12a expression plasmid was generated by deleting MBP from pDEST-hisMBP-AsCpf1-EC (Addgene: 79007 (*41*)), and transformed into Rosetta(DE3)pLysS Competent Cells (Novagen) for expression. AsCas12a protein was purified as previously described (*41*).

*ACTB and ACTG1 gene editing*: sgRNAs were generated using GeneArt Precision gRNA Synthesis Kit (Thermo Fisher Scientific: A29377). The sgRNA target sequences were:

*ACTB-Δ4nt^+/-^*: GCTATTCTCGCAGCTCACCA
*ACTB-insPTC^+/-^*: CCTGGGGCGCCCCACGATGG
*ACTG1-insPTC^+/-^*: CGAGCTGCGCGTGGCCCCGG

The corresponding single-stranded DNA (ssDNA) oligo repair templates were:

*ACTB-Δ4nt^+/-^*: gaagtggccagggcgggggcgacctcggctcacagcgcgcccggctattctcgcagctcaccatgatgatatcgccgcgctcgtcgtcgacaacggctccggcatgtgcaaggccggcttcgcgggcgacgatg
*ACTB-insPTC^+/-^*:
ctcccggggctgccccacccagccagctcccctacctggtgcctggggcgccccacgatggaAgg**TTATTA**gaagac ggcccggggggcatcgtcgcccgcgaagccggccttgcacatgccggagccgttgtcgacgacgag
*ACTG1-insPTC^+/-^*:
caccaactgggacgacatggagaagatctggcaccacaccttctacaacgagctgcgcgtggcc**TAATAA**ccggagga gcacccagtgctgctgaccgaggcccccctgaaccccaaggccaacagagagaag

Electroporation was conducted using the Neon Transfection System: 0.6 μg sgRNA was incubated with 3 μg HiFi Cas9 for 10 min at room temperature, and electroporated with 2 × 10⁵ H9 cells and ssDNA oligo.

To generate *ACTB-Δgene^+/-^* mutant, the sgRNA target sequences GGCGCCCTATAAAACCCAG and GAGGCCAAGTGTGACTTTG were inserted into the BbsI site in pPAX-HiFi and pU6-HiFi, respectively. pPAX-HiFi and pU6-HiFi were modified from SpCas9(BB)-2A-GFP (PX458) (Addgene: 48138 (*42*)) and pU6-(BbsI) CBh-Cas9-T2A-mCherry (Addgene: 64324 (*43*)) by introducing an R691A mutation into the SpCas9 coding sequence. H9 cells (1 × 10^6^) were transfected with 2.5 μg each of the two sgRNA-expressing plasmids. Cells expressing both GFP and mCherry were sorted as single cells into 96-well plates.

##### KINDLIN2 gene editing

To generate *KINDLIN2^+/-^*mutants, 80 pmol of Alt-R CRISPR-Cas12a crRNA targeting sequence GTGACAGTGCTTTGTCAGAAGGCA was incubated with 63 pmol of AsCas12a protein for 10 minutes at room temperature and electroporated into 2 × 10⁵ H9 cells along with 40 pmol of a ssDNA oligo: cacttatgatgctcatgatggaagccccttgtcaccaacttctgcttggtttggtgacagtgctttgtcagaa**TAATAA**ggcaatcctggt atacttgctgtcagtcaaccaatcacgtcaccagaaatcttggcaaaaatgttcaagcc and 39 pmol of Alt-R Cpf1 Electroporation Enhancer.

##### Genotyping (MiSeq)

Genomic DNA for genotyping was prepared by crude lysis. Briefly, culture medium was aspirated and 30 µL PBND buffer containing proteinase K (0.2 mg/mL; 50 mM KCl, 10 mM Tris-HCl pH 8.3, 2.5 mM MgCl₂, 0.45% NP-40, 0.45% Tween-20) was added to each well, incubated, and heat-inactivated to generate lysates.

Amplicon libraries were generated using a two-step PCR protocol. In the first PCR (PCR1), locus-specific primers (Primer 1 and Primer 2, see Supplementary Table S1) were used to amplify the genomic region surrounding the CRISPR target site. Primer 1 contained the Illumina Read 1 priming sequence (ACACTCTTTCCCTACACGACGCTCTTCCGATCT) fused to a gene-specific sequence (18–22 nt), and Primer 2 contained the Illumina Read 2 priming sequence (GTGACTGGAGTTCAGACGTGTGCTCTTCCGATCT) fused to the reverse-complement gene-specific sequence (18–22 nt). The region of interest was positioned within ∼100–130 bp of the primers so that the editing site was covered. PCR1 reactions (10 µL) contained 1 µL lysate, 2× NEBNext Ultra II Q5 Master Mix (NEB), and 0.07 µL of each primer (100 µM).

In the second PCR (PCR2, index PCR), Illumina adapters and sample indices were added using a universal P5 primer (Primer 3; AATGATACGGCGACCACCGAGATCTACACTCTTTCCCTACACGACGCTCTTCCGATCT) and a well-specific indexed P7 primer (Primer 4; CAAGCAGAAGACGGCATACGAGATNNNNNNNNGTGACTGGAGTTCAGACGTGTGCT, where NNNNNNNN is an 8-nt index). PCR2 reactions (10 µL) contained 1 µL PCR1 product, 2× NEBNext Ultra II Q5 Master Mix, 0.1 µL universal P5 primer (100 µM), and 1 µL indexed P7 primer (10 µM).

For pooling, 3 µL of each PCR2 product was combined, purified using the QIAquick PCR Purification Kit (Qiagen: 28104), and eluted in 40 µL EB. Library concentration was measured by NanoDrop and adjusted to 4 nM. Pooled libraries were sequenced on an Illumina MiSeq using MiSeq Reagent Nano Kit v2 (300-cycles; Illumina: MS-103-1001) with ∼10 pM loading and 10–15% PhiX spike-in; index reads were 8 nt, and read length/direction were chosen to cover the CRISPR target site. Sequence data were analyzed using OutKnocker (outknocker.org) for alignment and indel calling; clones were classified as wild-type or mutant based on the frequency and nature of editing events at the target locus. For heterozygous mutants, there should be both wild-type and mutant reads (typically 50:50).

#### Transgene insertion in hESCs and HEK293FT cells

*Plasmid construction for doxycycline-inducible piggyBac transgenes*: A human *ACTB* genomic fragment corresponding to hg38 chr7:5527048–5530601 (3562 bp; containing *ACTB* exons E1–E6 and intervening introns/UTRs as diagrammed in Fig. 2B) was synthesized (GenScript, Addgene: 255705) and cloned into a Gateway entry vector (pGenDONR, attL1/attL2-flanked). An *ACTB-insPTC* variant containing TAATAA (two adjacent stop codons) was generated from the wild-type entry plasmid by site-directed mutagenesis using the following primers ccgtcttc**TAATAA**ccctccatcgtggggcgccccaggcacc (forward) and atggaggg**TTATTA**gaagacggcccggggggcatcgtcgcccg (reverse), followed by Gibson assembly back into the same entry backbone (Addgene: 255706). Wild-type and *ACTB-insPTC* entry clones were transferred by Gateway LR recombination into a doxycycline-inducible piggyBac destination vector (Backbone: PB-TAG-ERP2, Addgene: 80479 (*44*); PB-TAG-ERP2-human-ACTB-WT, Addgene: 255707; PB-TAG-ERP2-human-ACTB-insPTC, Addgene: 255708). *ACTB-insPTC* transgene has the same TAATAA insertion location as used in endogenous *ACTB-insPTC* mutant hESC line (Fig. S1A).

##### Stable genomic integration of the transgene in hESCs

H9 hESCs were dissociated to single cells using StemPro® Accutase® Cell Dissociation Reagent (Thermo Fisher Scientific: A1110501). 50,000 cells were plated per well of a 12-well plate and transfected at the time of plating (cells in suspension prior to attachment) using Lipofectamine™ Stem Transfection Reagent (2 µL, Thermo Fisher Scientific: STEM00008) with 100 ng piggyBac transgene plasmid and 100 ng Super PiggyBac transposase plasmid (System Biosciences: PB210PA-1). ROCK inhibitor (Y-27632; 10 µM; STEMCELL™ Technologies: 72304) was included during plating/transfection. Medium was changed the next day. Puromycin selection was initiated ∼7 days post-transfection (0.25 µg/mL for the first 4 days, 0.5 µg/mL thereon) and maintained until near-complete GFP induction was observed upon doxycycline treatment (polyclonal stable populations). Transgene expression was induced with doxycycline (1 µg/mL; Sigma: D9891) for 24 h.

##### Stable genomic integration of the transgene in HEK293FT cells

HEK293FT cells were plated at 0.2×10^6^ cells/well (12-well plate) and transfected the next day using Lipofectamine™ 3000 Transfection Reagent (2 µL, Thermo Fisher Scientific: L3000008) with 500 ng piggyBac transgene plasmid and 500 ng Super PiggyBac transposase plasmid (System Biosciences PB210PA-1). Puromycin (1 µg/mL) was added three days after transfection and maintained until stable polyclonal populations were obtained. Transgene expression was induced with doxycycline (1 µg/mL; Sigma: D9891) for 24 h or 48 h.

#### Cre-loxP recombination in MKFs

To induce Cre-mediated recombination in MKFs, cells were plated at 0.4×10^6^ cells per well in a 6-well plate and transfected with 1 µg/well of a Cre recombinase plasmid expressing GFP (Addgene: 20781, Tyler Jacks lab) using Lipofectamine™ 3000 Transfection Reagent (5 µL/well). The next day, GFP-positive cells were single-cell sorted using flow cytometry into 96-well plates. Clones were expanded (e.g., to 48-well plates) prior to genotyping. Genotyping was performed approximately 2 weeks after transfection.

For genotyping, genomic DNA was extracted using the Monarch Genomic DNA Purification Kit (NEB: T3010S). The floxed *Kindlin2* locus was genotyped by PCR using 2× Phanta Flash Master Mix (Dye Plus) (Vazyme/Cellagen Technology: P520) with primers TCTCGAAACAGCAGTCAGAGG (forward) and CAAGCATCCACTGTAGACCGA (reverse). Expected PCR product sizes were 1501 bp (wild-type) and 827 bp (deletion/recombined allele). PCR products were resolved by agarose gel electrophoresis (Fig. S1C). The mutant (827 bp) band was gel-extracted and verified by Sanger sequencing. Only wild-type or heterozygous recombined, but no homozygous recombined, clones were recovered despite multiple attempts with Cre recombinase.

Following single-cell sorting, colony survival was assessed by the number of wells yielding viable colonies. 48 of 196 single cells survived after transfection with a control (non-Cre) plasmid, whereas 49 of 576 single cells survived after transfection with the Cre recombinase plasmid. Of the 49 Cre-transfected clones that were expanded and genotyped, 32/49 were heterozygous recombinants, 17/49 were wild-type, and 0/49 were homozygous recombinants. Under a simple independent-allele mutation model, the absence of homozygous *Kindlin2^−/−^* clones deviated significantly from expectation (chi-square goodness-of-fit, p < 0.001), suggesting that complete loss of *Kindlin2* is lethal under these conditions.

### RNA expression analyses

#### RNA purification

Total RNA was isolated using TRIzol Reagent (Thermo Fisher Scientific: 15596018) followed by column-based cleanup. For a 12-well plate, 300 µL TRIzol was added directly to each well, and lysates were transferred to 1.7-mL microcentrifuge tubes. Chloroform (60 µL) was added, tubes were vigorously shaken, incubated for ∼2 min at room temperature, and centrifuged at 12,000 × g for 15 min at 4°C. The aqueous phase was transferred to a new tube, mixed with an equal volume of 100% ethanol, and loaded onto RNA Clean & Concentrator-5 columns (Zymo Research: R1013), followed by purification according to the manufacturer’s protocol. Briefly, columns were centrifuged at >12,000 × g for 1 min at room temperature, washed with 400 µL RNA Prep Buffer, then 700 µL RNA Wash Buffer, followed by an additional wash with 400 µL RNA Wash Buffer. Columns were centrifuged for 2 min to dry the membrane, and RNA was eluted in 20 µL RNase-free water.

To remove residual genomic DNA, samples were treated with TURBO DNase (Thermo Fisher Scientific: AM2239) and re-purified. For each 20-µL RNA sample, 2 µL TURBO DNase, 3 µL 10× DNase buffer, and 5 µL nuclease-free water were added (30-µL total), and reactions were incubated at 37°C for 30 min. For cleanup, 70 µL water, 200 µL RNA Binding Buffer, and 300 µL 100% ethanol were added, and the mixture was applied to RNA Clean & Concentrator columns. Columns were processed as above (400 µL RNA Prep Buffer; 700 µL and 400 µL RNA Wash Buffer; final 2-min spin), and RNA was eluted in 15–20 µL RNase-free water. Purified RNA was quantified by NanoDrop spectrophotometry and stored at −80°C until use.

#### Reverse transcription and quantitative PCR (RT-qPCR)

For reverse transcription, 2 µg of total RNA was used for each +RT (with Reverse Transcriptase) and −RT (without Reverse Transcriptase) control reaction. cDNA synthesis was performed using SuperScript IV Reverse Transcriptase (Thermo Fisher Scientific: 18090050) following the manufacturer’s instructions, with minor changes. For each 20-µL reaction, RNA (2 µg in 12 µL), 10 mM dNTP mix (1 µL; NEB N0447S), and either 1 µL oligo(dT)₍₂₀₎ (50 µM; Invitrogen 18418020) and/or 1 µL random hexamers (50 µM; Invitrogen N8080127) were combined, heated to 65°C for 5 min, and placed on ice for at least 1 min. Then, 4 µL 5× SuperScript IV buffer, 1 µL 100 mM DTT, and 1 µL SuperScript IV were added (water was substituted for enzyme in −RT controls). Reactions were incubated at 23°C for 10 min, 50°C for 30 min, and 80°C for 10 min, then held at 4°C.

Resulting cDNA was diluted 1:10 by adding 180 µL nuclease-free water to each 20-µL reaction. qPCR was performed on an Applied Biosystems real-time PCR instrument using SYBR Green master mix and gene-specific primers (listed in Supplementary Data S1). All primers were validated by gel electrophoresis and melt-curve analysis. Typically, 2–2.5 µL of diluted cDNA was used per 10-µL qPCR reaction. No-RT controls were included for each RNA sample.

Relative mRNA expression was calculated using the 2^−ΔΔCt method, with *PGK1* or *GAPDH* as the reference gene, as indicated in figure legends. For comparison between two groups, ΔCt values (Ct_gene of interest − Ct_housekeeping) were used to perform two-tailed Student’s t-tests on biological replicates, and p-values < 0.05 were considered significant. Details of n and statistical tests used for each experiment are provided in the figure legends.

#### RNA-seq library preparation, sequencing, and data analysis

Total RNA integrity was assessed using an Agilent Bioanalyzer, and only samples with RNA integrity number (RIN) > 9 were used for library preparation. For each sample, 1 µg total RNA was used as input. Polyadenylated RNA was enriched using the NEBNext Poly(A) mRNA Magnetic Isolation Module (NEB: E7490S), and strand-specific libraries were prepared with the NEBNext Ultra II Directional RNA Library Prep Kit (NEB: E7765S) according to the manufacturer’s instructions. Libraries were PCR-amplified for 8 cycles, quantified, pooled, and multiplexed for sequencing on an Illumina HiSeq platform using standard paired-end protocols.

Raw reads were demultiplexed, and adapter sequences were removed using Cutadapt (v1.18) (*45*). For visualization on a genome browser, adapter-trimmed paired-end reads were aligned to the human reference genome (GRCh38/hg38) using HISAT2 (v2.1.0) (*46*) with default parameters and forward–reverse orientation (--fr). Resulting SAM files were converted to sorted, indexed BAM files using SAMtools (v1.9) (*47*). Genome-wide coverage tracks were generated from BAM files using deepTools bamCoverage (v3.5.6) (*48*), with counts normalized as counts per million mapped reads (CPM) and a bin size of 10 bp. Indexed BAM or bigWig coverage tracks were used for alignment visualization on the Integrative Genomics Viewer (IGV) (*49*).

For transcript-level quantification and differential expression analysis, we used kallisto and sleuth (*50, 51*). Adapter-trimmed reads were pseudoaligned with kallisto (v0.46.2) to an Ensembl GRCh38 transcriptome index (release 96). Quantification was run in paired-end mode with reverse-forward strand specificity (--rf-stranded), 4 threads, and 100 bootstrap samples.

Fragment length parameters (mean 138 bp, SD 20 bp) were estimated from read length distributions of the trimmed FASTQ files and supplied to kallisto (-l 138 -s 20). Transcript-level abundances and bootstrap estimates were imported into R and analyzed with sleuth for differential expression. Genes/transcripts with a false discovery rate (FDR, q-value) < 0.01 were considered significantly differentially expressed.

For allele-specific analysis at the *ACTB* locus, HISAT2-aligned BAM files were visualized in IGV. For a heterozygous SNP in the 3′ UTR, read counts supporting either the reference (G) or alternate (A) allele were obtained from the IGV coverage track, and allelic ratios were calculated per sample.

Raw FASTQ files, alignments in bigWig format, kallisto quantification outputs, and the detailed command-line workflows used for processing and analysis are available at GEO under accession GSE317480.

### siRNA knockdowns

Transient knockdown in hESCs was performed using siRNA transfection with Lipofectamine™ RNAiMAX Transfection Reagent (Thermo Fisher Scientific: 13778150) according to the manufacturer’s instructions. For transfection, hESCs were dissociated to single cells with Accutase® Cell Dissociation Reagent (6 mL per 10-cm dish, Thermo Fisher Scientific: A1110501), diluted with 14 mL E8 medium, and centrifuged at 150 × g for 3 min. Pelleted cells were resuspended in E8 medium and counted, then seeded into 12-well plates at 0.2 × 10⁶ cells/mL. ROCK inhibitor (Y-27632; 10 µM; STEMCELL™ Technologies: 72304) was included during plating/transfection. siRNA–RNAiMAX cocktails were prepared in Opti-MEM (Thermo Fisher Scientific: 31985062) to a final concentration of 50 nM siRNA and 2 µL RNAiMAX per well, and added immediately to the cells at the time of plating (before attachment). Knockdown was carried out for 48–52 h before harvest (Fig. 4A–C: 52 h; Fig. 4D, E: 48 h). Total RNA was then extracted, and target gene knockdown was assessed by RT-qPCR as described above.

The following siRNAs were used:

SRF: FlexiTube siRNA Hs_SRF_5 (Qiagen: SI02757622)
MRTF-A (MKL1): ON-TARGETplus MKL1 (Horizon Discovery: L-015434-00-0005)
UPF1: ON-TARGETplus UPF1 (Horizon Discovery: J-011763-07-0005)
SMG6: ON-TARGETplus SMG6 (Horizon Discovery: J-017845-09-0005)
UPF3A: ON-TARGETplus UPF3A (Horizon Discovery: J-012872-10-0010)
ILF3: ON-TARGETplus ILF3 (Horizon Discovery: L2-012442-01-0005)
ON-TARGETplus Non-targeting Control Pool (Horizon Discovery: D-001810-10-05; used in Fig. 4A–C)
ON-TARGETplus 2.0 Non-targeting Control Pool (Horizon Discovery: D2-001810-10-05; used in Fig. 4D, E)

### Immunofluorescence staining, image acquisition, and analysis

#### Immunostaining

Actin staining was performed following the protocol available on the Mitchison Lab website (https://mitchison.hms.harvard.edu/resources: “Fluorescence Procedures for the Actin and Tubulin Cytoskeleton in Fixed Cells”) with minor modifications. Cells were cultured on glass-bottom chamber slides, rinsed once in PBS, and fixed in 4% formaldehyde in PBS for 13 min at room temperature. After three washes in PBS, cells were permeabilized in 0.1% Triton X-100 in PBS for 5 min, then rinsed three times in 0.1% Triton X-100/PBS. F-actin and G-actin were labeled by incubating cells for 15 min at room temperature in PBS containing 0.1% BSA, Alexa Fluor 488 Phalloidin (1:400; Thermo Fisher: A12379), and Alexa Fluor 594–conjugated DNase I (1:500; Thermo Fisher: D12372). Samples were washed three times in PBS, counterstained with DAPI (1 µg/mL in PBS, 5 min), and mounted in VECTASHIELD Antifade Mounting Medium (Vector Laboratories: H-1000-10).

#### Image acquisition

Images were acquired on a Nikon Ti2 inverted microscope equipped with a Yokogawa CSU spinning disk confocal unit, a Nikon Plan Apo λ 60×/1.40 NA oil-immersion objective (MRD01605), a Hamamatsu ORCA-Fusion sCMOS camera (C14440-20UP), a Lumencor Sola II LED illuminator, and a Nikon LUN-F multi-laser launch, controlled by NIS-Elements AR software. Z-stacks were collected with a voxel size of 0.108 × 0.108 × 0.3 µm³.

#### Quantifying fluorescence

For quantitative analysis of F-actin and G-actin, image processing and measurement were performed in Python (v3.x). Analysis code is available on Github (https://github.com/harleensaini/MoazedLab_hESC-actin-quantification) and raw imaging data are available on Zenodo (https://doi.org/10.5281/zenodo.19556498). For each ND2 stack, raw data were first background-subtracted on a per-channel basis by subtracting the minimum intensity value of each channel across the 3D stack. For quantification, 3D stacks were collapsed along the z-axis by summing intensities for the G-actin (DNase I) and F-actin (phalloidin) channels separately. To define the cell area within each field of view, an actin-based mask was generated from the phalloidin and DNase I channels. Briefly, background-subtracted phalloidin and DNase I channels were summed along z to produce a 2D actin intensity image, followed by histogram equalization, Gaussian blurring, and global thresholding using the triangle method.

The resulting binary image was refined by hole filling to generate a contiguous mask corresponding to the area occupied by cells. Masks were visually inspected for every image to verify accurate delineation of cell regions. Within the actin mask for each image, total fluorescence intensities of the summed-z G-actin and F-actin images were computed and normalized by the number of mask pixels to yield an average intensity per unit area. Each data point shown in Figure 1F corresponds to a single image (field of view), encompassing many cells. Multiple independent images were analyzed per genotype.

### Western blots

Cells were lysed directly in 1× Laemmli sample buffer (Bio-Rad 1610747) and briefly vortexed. Lysates were heated at 95°C for 5 min and separated by SDS–PAGE on 10% Mini-PROTEAN TGX precast gels (Bio-Rad) or homemade 10% SDS-PAGE gels. Proteins were transferred to PVDF membranes using the Trans-Blot Turbo transfer system (Bio-Rad). Membranes were blocked in 5% (w/v) nonfat dry milk in TBST (TBS, 0.1% Tween-20) for at least 1 h at room temperature, incubated with primary antibodies overnight at 4°C, washed in TBST (3 × 10 min), and then incubated with HRP-conjugated secondary antibodies for 1 h at room temperature, followed by TBST washes (3 × 10 min). Signals were detected using SuperSignal West Pico PLUS Chemiluminescent Substrate or SuperSignal West Femto Maximum Sensitivity Substrate (Thermo Fisher Scientific). Protein loading and transfer were assessed by Ponceau S staining of membranes prior to blocking and/or by post-imaging Coomassie staining with SimplyBlue SafeStain (Thermo Fisher). Housekeeping proteins (e.g., GAPDH) were used as loading controls.

Primary antibodies and dilutions:

β-actin (ACTB): mouse monoclonal, Abcam: ab6276; 1:5000
SRF: mouse monoclonal, Proteintech: 66742-1-Ig; 1:1000
MRTF-A (MKL1): rabbit polyclonal, Thermo Fisher: PA5-56557; 1:500
UPF1: rabbit monoclonal, Abcam: ab109363; 1:5000
UPF3A: rabbit polyclonal, Abclonal: A15893; 1:1000
SMG6: rabbit monoclonal, Abcam: ab87539; 1:1000
ILF3: rabbit monoclonal, Abcam: ab92355; 1:3000
GAPDH: rabbit monoclonal, Abcam: ab125247; 1:2000

Secondary antibodies (1:5000 dilution):

Rabbit TrueBlot ULTRA: Anti-Rabbit IgG HRP (Rockland: 18-8816-31),
Mouse TrueBlot ULTRA: Anti-Mouse IgG HRP (Rockland: 18-8817-31).

### Re-plotting published flow-FISH screen data (El-Brolosy et al., 2026)

Flow-FISH CRISPR screen results were obtained from Supplementary Table S3 of El-Brolosy et al. (*36*). We used the gene-level MAGeCK summary statistics provided by the authors for the *Actg1-NSD* (nonstop decay) screen and the matched wild-type (WT) counter-screen. In the original study, cells were stained by flow-FISH for *Actg2* and a housekeeping transcript (*Rpl13a*) and then sorted based on the *Actg2*:*housekeeping* fluorescence ratio into *Actg2*-high and *Actg2*-low fractions prior to sgRNA sequencing and MAGeCK analysis.

We imported the “Actg1 NSD screen” and “WT counter screen” sheets from Table S3 (El-Brolosy et al., 2026) into R (RStudio) and used the reported MAGeCK outputs. Genes were classified as *Actg2*-up hits (pos|lfc > 0.5, FDR < 0.05), *Actg2*-down hits (pos|lfc < −0.5, FDR < 0.05), or not significant. For each gene, we assigned a p-value and FDR by selecting the positive-selection values (pos|p-value, pos|fdr) when pos|lfc > 0 and the negative-selection values (neg|p-value, neg|fdr) when pos|lfc < 0, and calculated −log10(p-value) for plotting. Volcano plots were generated with ggplot2 (*52*), highlighting a predefined list of genes of interest (*Ilf3*, *Mkl1*, *Srf*, *Actg1*, *Actb*, *Acta1*, and *Actg2*). No new statistical tests were applied to the screen data; all values are taken directly from the published MAGeCK analysis (*36*).

The rescued *Actg1-NSD* screen was performed in duplicate, whereas the WT counter-screen was performed as a single replicate in the original study (*36*). Accordingly, differences in statistical significance between the two screens should be interpreted cautiously, as they may reflect differences in replication and statistical power in addition to biological effects.

## Supplementary Materials

**Other Supplementary Materials for this manuscript include the following:**

Data S1 (Excel file): Primer sequences for genotyping and RT-qPCR.

Data S2 (Excel file): Ct values and calculations for RT-qPCR data in Fig. 1.

Data S3 (Excel file): Ct values and calculations for RT-qPCR data in Fig. 2.

Data S4 (Excel file): Ct values and calculations for RT-qPCR data in Fig. 4.

Data S5 (Excel file): Ct values and calculations for RT-qPCR data in Fig. S5

Data S6 (PDF): Uncropped western blot images corresponding to all main and supplementary figures.

**Fig. S1.**
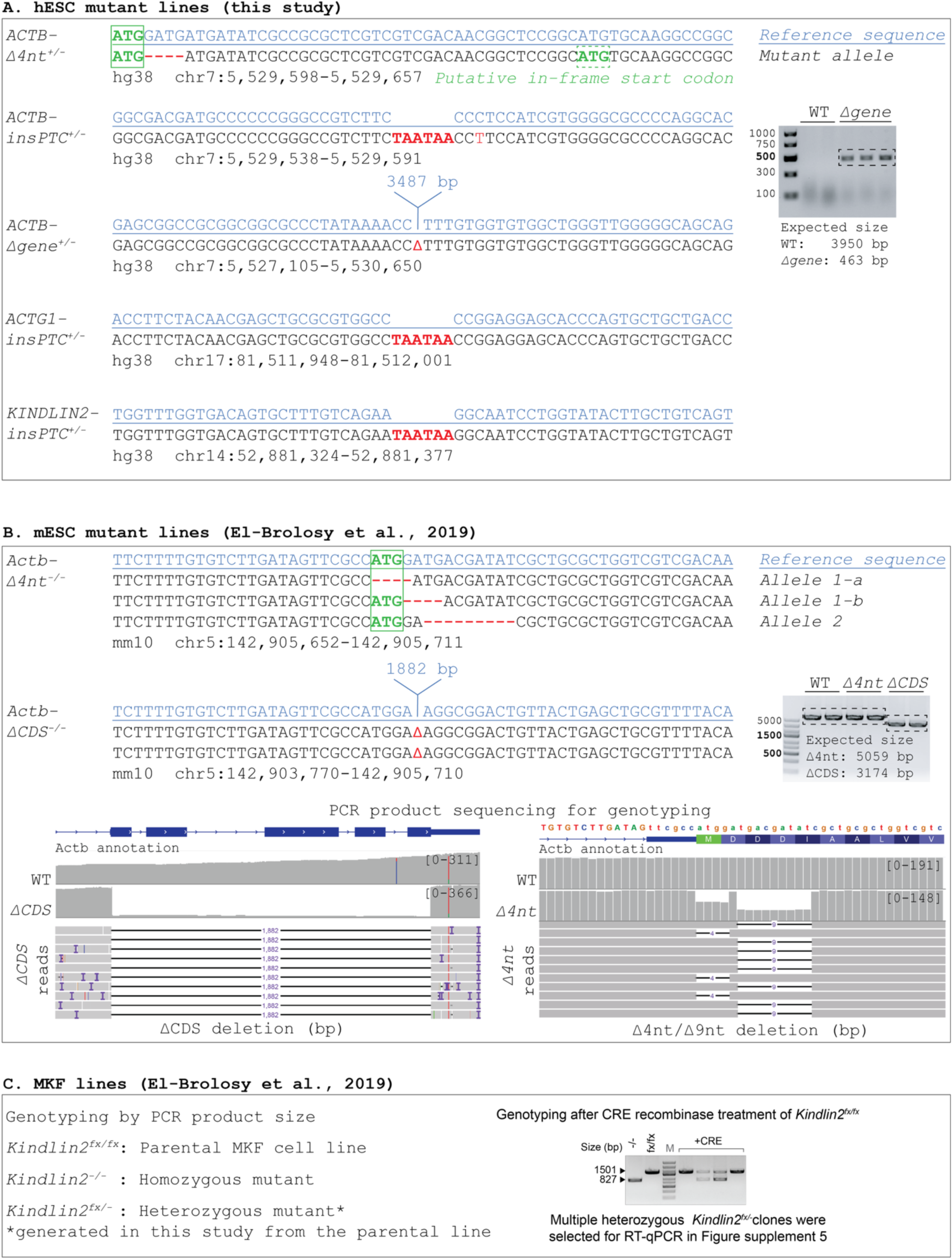
Genotyping human and mouse cell lines. **(A)** hESC mutant lines: For each hESC genotype, the reference sequence and the corresponding mutant allele sequence are shown across the edited region; mutations (insertions/deletions) are shown in red font, translation start codons are enclosed in green box. Genomic coordinates (hg38) denote the reference sequence interval shown for each alignment. Inset: PCR-based genotyping of the *ACTB-Δgene* allele resolved by agarose gel electrophoresis, with mutant allele showing smaller PCR product size. **(B)** mESC mutant lines: mESC mutant genotypes are shown with reference and mutant sequences, along with a representative PCR genotyping gel. For each homozygous mutant line, both Allele 1 and Allele 2 sequences are displayed, with the corresponding chromosomal coordinates (mm10) indicated as in (A). For *Actb-Δ4nt^-/-^*, Allele 1-a and 1-b show two possible alignments for the same sequence. Bottom: Integrative Genomics Viewer (IGV) snapshots of long-read amplicon sequencing across the target locus, showing read depth (y-axis scaled 0–375 reads) and individual read alignments confirming the indicated mutant alleles. Amplicons were sequenced using Oxford Nanopore Technologies, and reads were aligned to the mouse reference genome (mm10) with minimap2 (default settings). **(C)** MKF mutant lines: MKF mutant genotypes are listed, with PCR-based genotyping of the *Kindlin2* locus shown by agarose gel electrophoresis, indicating wild-type and recombined (deletion) alleles.

**Fig. S2.**
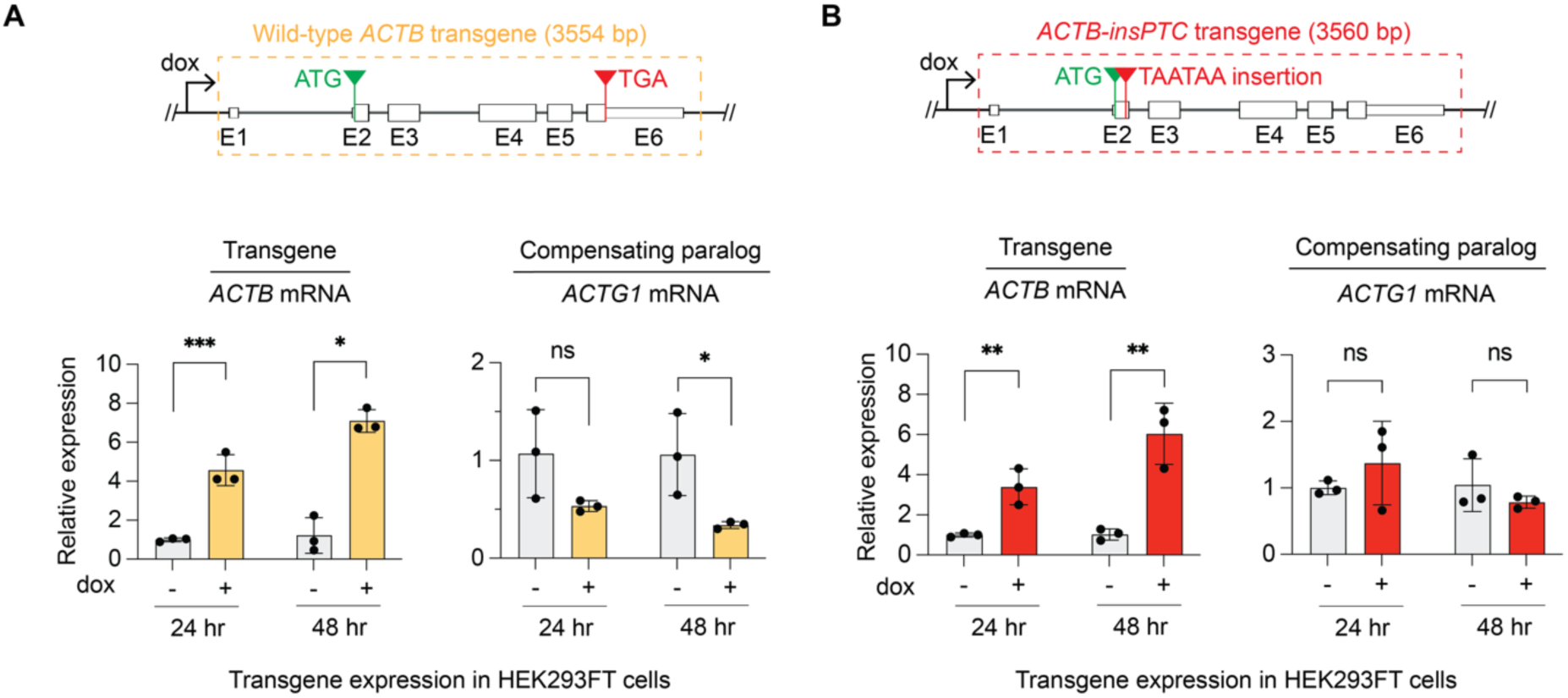
Transgenic expression of mutant ACTB mRNA does not induce genetic compensation in HEK293FT cells. **(A, B)** RT-qPCR analysis *ACTB* and *ACTG1* after 24 or 48 hours of wild-type (A) or insPTC (B) *ACTB* transgene induction by doxycycline. Relative expression was calculated using the 2^-ΔΔCt method, normalized to *PGK1* mRNA.

**Fig. S3.**
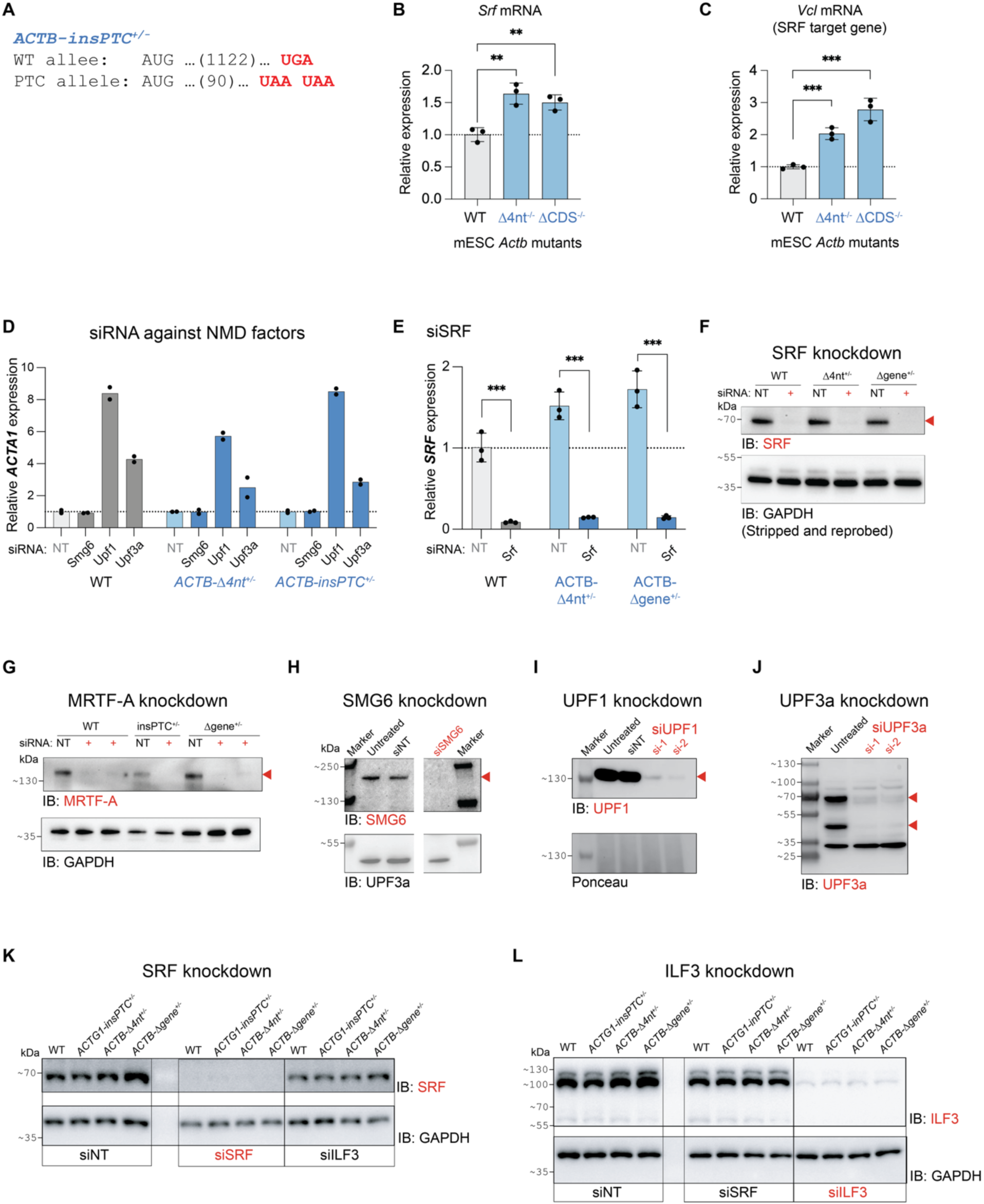
Validation of knockdowns in hESCs and SRF pathway activation in mESC *Actb* mutants. **(A)** Sequence showing location of PTCs in *ACTB-insPTC^+/–^* allele compared to wild-type allele. **(B)** RT-qPCR analysis of *Srf* mRNA expression in wild-type mESCs and *Actb* mutant mESCs. **(C)** RT-qPCR analysis of *Vcl* mRNA expression in wild-type mESCs and *Actb* mutant mESCs. **(D)** RT-qPCR analysis of *ACTA1* mRNA expression after siRNA-mediated knockdown of NMD factors (SMG6, UPF1, UPF3A), displayed with an alternative normalization: for each genotype, expression values are normalized to the corresponding siNT control to emphasize differences between siNT and siNMD within each genotype. In contrast, Fig. 4C shows the same data normalized to the siNT condition in wild-type cells. **(E)** Validation of *SRF* knockdown in hESCs by RT-qPCR. **(F-L)** Knockdown validation by immunoblotting. **(F)** SRF knockdown corresponding to Fig. 4A. **(G)** MRTF-A knockdown corresponding to Fig. 4B. **(H–J)** SMG6, UPF1, and UPF3A knockdowns, respectively, corresponding to Fig. 4C. **(K, L)** SRF and ILF3 knockdowns, respectively, corresponding to Fig. 4D and 4E.

**Fig. S4.**
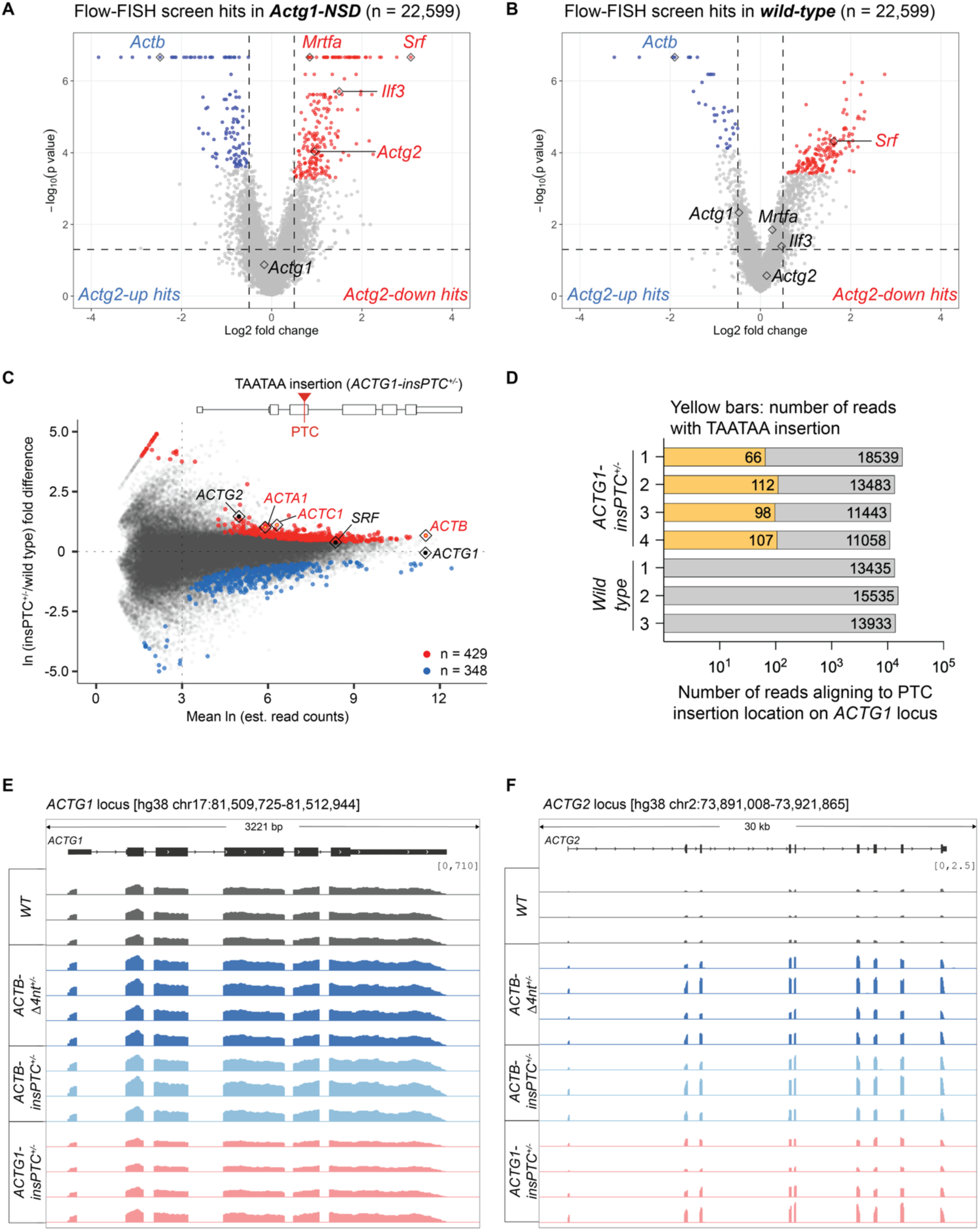
Re-plotted flow-FISH CRISPR screen and *ACTG1-insPTC* RNA-seq analyses. **(A, B)** Published genome-wide CRISPR flow-FISH screens identifying regulators of *Actg2* expression (El-Brolosy et al., 2026; Table S3), highlighting the requirement for Srf, Mrtfa, and other factors in addition to Ilf3. Volcano plots show gene-level MAGeCK results for (A) the *Actg1-NSD* (nonstop decay) mutant line and (B) the matched wild-type (WT) counter-screen. Each point represents a gene targeted by a pool of sgRNAs; the x-axis shows log2 fold change (pos|lfc), and the y-axis shows −log10(p-value), from the MAGeCK output (Table S3). The dashed horizontal line denotes p = 0.05. Genes were classified as “*Actg2*-up hits” (red; pos|lfc > 0.5, FDR < 0.05) or “*Actg2*-down hits” (blue; pos|lfc < −0.5, FDR < 0.05); all others are shown in grey. Selected genes of interest (*Actb*, *Actg1*, *Actg2*, *Srf*, *Mkl1* (*Mrtfa*), *Ilf3*) are highlighted. Among the highlighted genes, *Srf* scored as an *Actg2*-up hit in both the WT counter-screen and the *Actg1-NSD* screen, consistent with its broad role in transcriptional regulation. In contrast, *Mkl1/Mrtfa* met the significance threshold in the *Actg1-NSD* screen but not in the WT counter-screen, suggesting that factors with general roles in actin/SRF signaling can score differently across screening contexts. **(C)** Scatter plot showing RNA-seq quantification, comparing log_e_ mean read counts and log_e_ fold difference in expression in *ACTG1-insPTC^+/–^*mutant versus wild-type hESCs per annotated mRNA isoform, plotted as in Fig. 3. Transcripts with significant changes in abundance (q-value < 0.01) are labeled in red (increased expression) or blue (decreased expression), all others are labeled in black. Actin paralogs and *SRF* transcription factor are annotated. A schematic of the *ACTG1-insPTC* mutant allele is shown above the plot. **(D)** Bar plot showing the total number of RNA-seq reads overlapping the TAATAA insertion site at the *ACTG1* locus (log_10_ scale). For each sample, the total bar length represents all reads mapping across the insertion site, and the yellow segment indicates the subset of reads that contain the TAATAA insertion (two adjacent stop codons) rather than the wild-type sequence. The very low number of reads bearing the insertion at steady state indicates efficient degradation of the mutant *ACTG1* transcript in *ACTG1-insPTC^+/-^*. **(E, F)** IGV screenshots of bigWig RNA-seq coverage across the *ACTG1* (E) and *ACTG2* (F) loci in WT and actin-mutant hESCs. Coverage is shown in CPM (counts per million) units, and tracks are displayed with group scaling. Each track represents an independent RNA-seq sample. (independent clones). Increased *ACTG2* expression in *ACTB* mutants is evident from elevated read coverage over the *ACTG2* gene body compared to WT; however *ACTG1-insPTC* mutants show clonal variability in *ACTG2* upregulation.

**Fig. S5.**
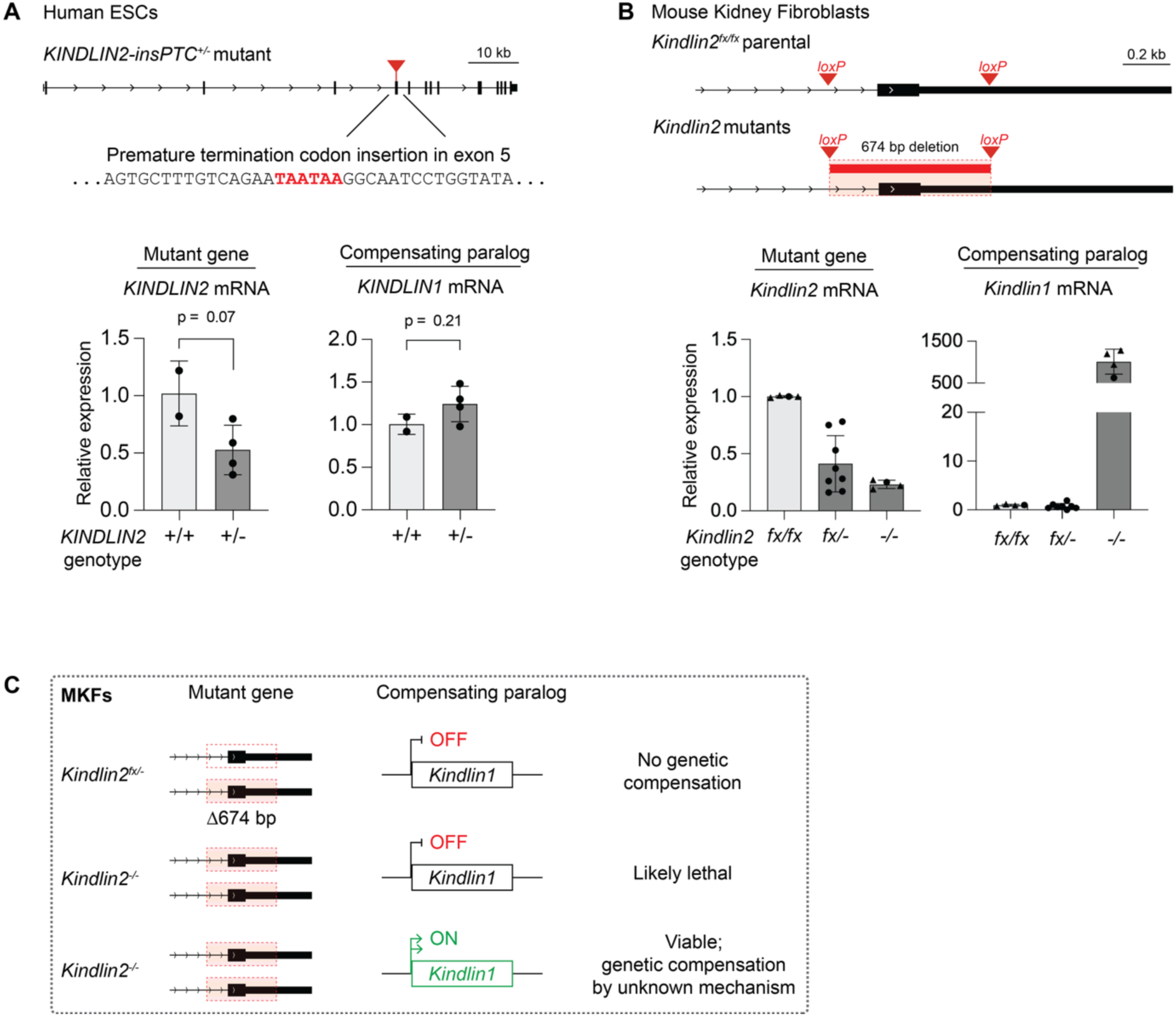
Heterozygous *Kindlin2* mutations do not trigger genetic compensation in hESCs and MKFs. **(A)** hESC *KINDLIN2* mutants. (Top) Schematic of the *KINDLIN2* locus showing the introduction of premature translation termination codons in exon 5 in human ESCs. (Bottom) RT-qPCR analysis of *KINDLIN2* and *KINDLIN1* mRNA expression in wild-type (+/+) and heterozygous (+/−) *KINDLIN2* mutant hESCs. Relative expression was calculated using the 2^−ΔΔCt method, normalized to *PGK1* mRNA and wild-type cells. Error bars denote standard deviation; p-values were calculated using two-tailed Student’s t-tests on ΔCt values (gene-of-interest – PGK1). **(B)** MKF *Kindlin2* mutants. (Top) Schematic of parental and mutant *Kindlin2* mouse kidney fibroblast (MKF) lines. The deleted segment in the homozygous *Kindlin2^−/−^* line is indicated. (Bottom) RT-qPCR analysis of *Kindlin2* and *Kindlin1* mRNA expression in parental (fx/fx), heterozygous (fx/−), and homozygous (−/−) *Kindlin2* MKFs. Expression was normalized to *Gapdh* and calculated as in (A). Fold changes from two independent experiments are shown, with data points from each experiment distinguished by circles or triangles. Error bars and p-values were calculated as in (A). **(C)** Model: Homozygous *Kindlin2* mutant MKFs may require *Kindlin1* expression and/or additional adaptive changes for viability, whereas heterozygous mutants do not exhibit compensatory upregulation of *Kindlin1*.

## Notes

### Competing Interest Statement

The authors have declared no competing interest.

## References

1. Z. Gu et al., Role of duplicate genes in genetic robustness against null mutations. Nature 421, 63–66 (2003).

2. Z. Kontarakis, D. Y. R. Stainier, Genetics in Light of Transcriptional Adaptation. Trends Genet 36, 926–935 (2020).

3. Y. Cho, Y. K. Kim, Multilayered regulation of cytoskeletal protein abundance: autoregulatory mechanisms of actin and tubulin. Exp Mol Med 58, 59–72 (2026).

4. C. W. Yeh et al., Altered assembly paths mitigate interference among paralogous complexes. Nat Commun 15, 7169 (2024).

5. A. Rossi et al., Genetic compensation induced by deleterious mutations but not gene knockdowns. Nature 524, 230–233 (2015).

6. M. A. El-Brolosy et al., Genetic compensation triggered by mutant mRNA degradation. Nature 568, 193–197 (2019).

7. Z. Ma et al., PTC-bearing mRNA elicits a genetic compensation response via Upf3a and COMPASS components. Nature 568, 259–263 (2019).

8. V. Serobyan et al., Transcriptional adaptation in Caenorhabditis elegans. Elife 9, (2020).

9. Z. Jiang et al., Parental mutations influence wild-type offspring via transcriptional adaptation. Sci Adv 8, eabj2029 (2022).

10. L. Falcucci et al., Transcriptional adaptation upregulates utrophin in Duchenne muscular dystrophy. Nature 639, 493–502 (2025).

11. M. F. Wilkinson, Genetic paradox explained by nonsense. Nature 568, 179–180 (2019).

12. A. Xie et al., Upf3a but not Upf1 mediates the genetic compensation response induced by leg1 deleterious mutations in an H3K4me3-independent manner. Cell Discov 9, 63 (2023).

13. J. M. Welker, V. Serobyan, E. Zaker Esfahani, D. Y. R. Stainier, Partial sequence identity in a 25-nucleotide long element is sufficient for transcriptional adaptation in the Caenorhabditis elegans act-5/act-3 model. PLoS Genet 19, e1010806 (2023).

14. L. Xie et al., Induction of a transcriptional adaptation response by RNA destabilization events. EMBO Rep 26, 2262–2279 (2025).

15. L. Falcucci, B. Juvik, D. Y. Stainier, Transcriptional adaptation: where mRNA decay meets genetic compensation. Curr Opin Genet Dev 93, 102369 (2025).

16. T. D. Pollard, Actin and Actin-Binding Proteins. Cold Spring Harb Perspect Biol 8, (2016).

17. R. Dominguez, K. C. Holmes, Actin structure and function. Annu Rev Biophys 40, 169–186 (2011).

18. K. Miyamoto, J. B. Gurdon, Transcriptional regulation and nuclear reprogramming: roles of nuclear actin and actin-binding proteins. Cell Mol Life Sci 70, 3289–3302 (2013).

19. C. Ampe, M. Van Troys, Mammalian Actins: Isoform-Specific Functions and Diseases. Handb Exp Pharmacol 235, 1–37 (2017).

20. A. S. Kashina, Regulation of actin isoforms in cellular and developmental processes. Semin Cell Dev Biol 102, 113–121 (2020).

21. P. Vedula, A. Kashina, The makings of the ‘actin code’: regulation of actin’s biological function at the amino acid and nucleotide level. J Cell Sci 131, (2018).

22. T. M. Bunnell, B. J. Burbach, Y. Shimizu, J. M. Ervasti, beta-Actin specifically controls cell growth, migration, and the G-actin pool. Mol Biol Cell 22, 4047–4058 (2011).

23. T. M. Bunnell, J. M. Ervasti, Delayed embryonic development and impaired cell growth and survival in Actg1 null mice. Cytoskeleton (Hoboken) 67, 564–572 (2010).

24. D. Tondeleir et al., Cells lacking beta-actin are genetically reprogrammed and maintain conditional migratory capacity. Mol Cell Proteomics 11, 255–271 (2012).

25. N. Malek et al., Knockout of ACTB and ACTG1 with CRISPR/Cas9(D10A) Technique Shows that Non-Muscle beta and gamma Actin Are Not Equal in Relation to Human Melanoma Cells’ Motility and Focal Adhesion Formation. Int J Mol Sci 21, (2020).

26. T. Ikeda et al., Srf destabilizes cellular identity by suppressing cell-type-specific gene expression programs. Nat Commun 9, 1387 (2018).

27. P. Vedula et al., Diverse functions of homologous actin isoforms are defined by their nucleotide, rather than their amino acid sequence. Elife 6, (2017).

28. S. Mouilleron, C. A. Langer, S. Guettler, N. Q. McDonald, R. Treisman, Structure of a pentavalent G-actin*MRTF-A complex reveals how G-actin controls nucleocytoplasmic shuttling of a transcriptional coactivator. Sci Signal 4, ra40 (2011).

29. D. Gau, P. Roy, SRF’ing and SAP’ing - the role of MRTF proteins in cell migration. J Cell Sci 131, (2018).

30. F. Miralles, G. Posern, A. I. Zaromytidou, R. Treisman, Actin dynamics control SRF activity by regulation of its coactivator MAL. Cell 113, 329–342 (2003).

31. G. Posern, A. Sotiropoulos, R. Treisman, Mutant actins demonstrate a role for unpolymerized actin in control of transcription by serum response factor. Mol Biol Cell 13, 4167–4178 (2002).

32. G. Posern, F. Miralles, S. Guettler, R. Treisman, Mutant actins that stabilise F-actin use distinct mechanisms to activate the SRF coactivator MAL. EMBO J 23, 3973–3983 (2004).

33. E. N. Olson, A. Nordheim, Linking actin dynamics and gene transcription to drive cellular motile functions. Nat Rev Mol Cell Biol 11, 353–365 (2010).

34. F. Parker, T. G. Baboolal, M. Peckham, Actin Mutations and Their Role in Disease. Int J Mol Sci 21, (2020).

35. E. P. Consortium, An integrated encyclopedia of DNA elements in the human genome. Nature 489, 57–74 (2012).

36. M. A. El-Brolosy et al., Mechanisms linking cytoplasmic decay of translation-defective mRNA to transcriptional adaptation. Science 391, eaea1272 (2026).

37. C. H. Lou et al., Nonsense-Mediated RNA Decay Influences Human Embryonic Stem Cell Fate. Stem Cell Reports 6, 844–857 (2016).

38. G. C. Pipes, E. E. Creemers, E. N. Olson, The myocardin family of transcriptional coactivators: versatile regulators of cell growth, migration, and myogenesis. Genes Dev 20, 1545–1556 (2006).

39. J. A. Zuris et al., Cationic lipid-mediated delivery of proteins enables efficient protein-based genome editing in vitro and in vivo. Nat Biotechnol 33, 73–80 (2015).

40. C. A. Vakulskas et al., A high-fidelity Cas9 mutant delivered as a ribonucleoprotein complex enables efficient gene editing in human hematopoietic stem and progenitor cells. Nat Med 24, 1216–1224 (2018).

41. J. K. Hur et al., Targeted mutagenesis in mice by electroporation of Cpf1 ribonucleoproteins. Nat Biotechnol 34, 807–808 (2016).

42. F. A. Ran et al., Genome engineering using the CRISPR-Cas9 system. Nat Protoc 8, 2281–2308 (2013).

43. V. T. Chu et al., Increasing the efficiency of homology-directed repair for CRISPR-Cas9-induced precise gene editing in mammalian cells. Nat Biotechnol 33, 543–548 (2015).

44. S. I. Kim et al., Inducible Transgene Expression in Human iPS Cells Using Versatile All-in-One piggyBac Transposons. Methods Mol Biol 1357, 111–131 (2016).

45. M. Martin, Cutadapt removes adapter sequences from high-throughput sequencing reads. EMBnet.journal 17, 10–12 (2011).

46. D. Kim, J. M. Paggi, C. Park, C. Bennett, S. L. Salzberg, Graph-based genome alignment and genotyping with HISAT2 and HISAT-genotype. Nat Biotechnol 37, 907–915 (2019).

47. P. Danecek et al., Twelve years of SAMtools and BCFtools. Gigascience 10, (2021).

48. F. Ramirez et al., deepTools2: a next generation web server for deep-sequencing data analysis. Nucleic Acids Res 44, W160–165 (2016).

49. J. T. Robinson et al., Integrative genomics viewer. Nat Biotechnol 29, 24–26 (2011).

50. H. Pimentel, N. L. Bray, S. Puente, P. Melsted, L. Pachter, Differential analysis of RNA-seq incorporating quantification uncertainty. Nat Methods 14, 687–690 (2017).

51. N. L. Bray, H. Pimentel, P. Melsted, L. Pachter, Near-optimal probabilistic RNA-seq quantification. Nat Biotechnol 34, 525–527 (2016).

52. H. Wickham, ggplot2: Elegant Graphics for Data Analysis. (Springer-Verlag, New York, 2016).

